# Decoding ATG9A Variation: A Comprehensive Structural Investigation of All Missense Variants

**DOI:** 10.64898/2026.02.03.703515

**Authors:** Mattia Utichi, Henri-Baptiste Marjault, Matteo Tiberti, Elena Papaleo

## Abstract

Macroautophagy (hereafter autophagy) is a cellular recycling pathway that requires different ATG (autophagy-related) proteins to generate double-membraned autophagosomes. ATG9A, a multi-spanning membrane protein, plays a crucial role in this process as the only transmembrane component of the core autophagy machinery. ATG9A functions as a lipid scramblase, redistributing lipids between membrane leaflets for the expanding autophagosome membrane. Structural studies have revealed that ATG9A forms a homotrimer with an interlocked domain-swapped architecture and a network of internal hydrophilic cavities. This configuration underlies its role in lipid transfer and membrane remodeling together with the lipid transporter ATG2A. ATG9A dysfunction has also been linked to human disease, as specific ATG9A mutations cause neurodevelopmental or neurodegenerative phenotypes. Additionally, ATG9A is altered in cancer, promoting pro-tumorigenic traits. However, most missense variants in ATG9A remain uncharacterized, posing a significant challenge for interpreting genomic data. In this study, we employed *in silico* saturation mutagenesis approach using the MAVISp (Multi-layered Assessment of VarIants by Structure) framework to predict the impact of every missense mutation in ATG9A. By analyzing multiple structural assemblies of ATG9A (monomer, trimer, and the ATG9A-ATG2A complex), we evaluated diverse mechanistic indicators of variant impact, including protein stability, long-range conformational changes, effects on multimerization interfaces, and alterations in post-translational modifications. We integrated the structure-based predictions with Variant Effect Predictors from recent deep-learning or evolutionary-based models and cross-referenced known variants catalogued in ClinVar, COSMIC, and cBioPortal. Finally, we predicted mechanistic indicators for all possible variants with structural coverage not yet reported in the disease-related databases supported by MAVISp. Our analyses identified a group of potentially damaging variants in ATG9A and the possible molecular mechanisms underlying their effects. Together, this work provides a roadmap for interpreting missense variants in autophagy regulators and highlights specific ATG9A mutations that deserve further investigation in the context of human disease.

## Introduction

Autophagy is a conserved catabolic process for degradation and recycling of cellular components. The formation of autophagosomes, the characteristic double-membraned vesicles of autophagy, requires the action of different ATG (Autophagy-related) proteins. Among these, ATG9A is the only transmembrane component of the core autophagy machinery and plays an essential role in autophagosome biogenesis^1^. Functionally, ATG9A acts as a lipid scramblase, facilitating the transbilayer movement of lipids between membrane leaflets to promote phagophore (nascent autophagosome) expansion^2^. Consistent with this activity, ATG9A dynamically traffics to sites of autophagosome initiation, contributing to membrane supply and remodeling during autophagosome formation^3^. ATG9A function depends critically on its continuous cycling between the trans-Golgi network, endosomes, and autophagosome formation sites, a process regulated by vesicular trafficking factors such as ULK1, AP-1/AP-4 complexes, and Rab GTPases^4–6^. Beyond its canonical role in autophagy, ATG9A also supports cellular homeostasis in other contexts, including maintaining neuronal integrity during development and regulating membrane dynamics under stress^7,8^. Dysregulation of ATG9A has been implicated in human disease: loss-of-function ATG9 mutations have been reported in patients with severe neurodevelopmental or neurodegenerative disorders^9–12^. Aberrant ATG9A expression or mislocalization is observed in several cancers^13,14^. These observations suggest that ATG9A plays an important role in human physiology and disease and raise the question of how changes in its sequence may affect autophagy and cellular homeostasis.

Recent advances in structural biology are enabling a deeper understanding of the molecular mechanisms of ATG9A. High-resolution cryo-EM structures of human ATG9A revealed that the protein assembles into a domain-swapped homotrimer, in which each protomer contributes several interlocking transmembrane helices to form a stable trimeric scaffold^2,15,16^. Interestingly, these structures revealed a broad system of internal cavities along with a central hydrophilic pore that runs through the membrane bilayer. The central pore, sealed on the cytosolic side by the C-terminal domain, can laterally accommodate lipid headgroups, providing a physical conduit for lipid flip-flop between leaflets^2,15^. This architecture supports ATG9A scramblase activity, which promotes lipid redistribution and supports phagophore expansion during autophagosome biogenesis^2,17^. Functional studies have confirmed the importance of ATG9A’s structural properties. For example, pore-blocking point mutations such as R422W abolish ATG9A’s ability to rescue autophagosome formation in ATG9A-deficient cells, demonstrating that pore integrity is essential for autophagy^16^. In addition to self-oligomerizing, ATG9A forms critical protein–protein interactions. Notably, ATG9A binds directly to ATG2A, a lipid-transfer protein, creating a functional complex that shuttles lipids from ATG2A to the expanding phagophore membrane^18,19^. This interaction allows ATG9A to supply lipids into the growing phagophore, a process required for autophagosome maturation. Computational and biophysical analyses further suggest that ATG9A homotrimers can induce membrane curvature: molecular dynamics simulations show that ATG9A induces dome-like curvature in the surrounding bilayer, consistent with its function as a membrane-shaping factor during phagophore growth^16^.

Given ATG9A central role in autophagy, perturbations to its structure may have significant functional consequences. Indeed, specific missense mutations in ATG9A have been reported to affect autophagy^16,20,21^, yet relatively few of the possible ATG9A variants have been experimentally characterized to date. Large-scale sequencing studies identify numerous rare variants in ATG9A in disease contexts. For example, cancer genomics resources such as COSMIC and cBioPortal catalog somatic *ATG9A* mutations across tumor types^22,23^, and ClinVar contains germline *ATG9A* variants observed in patients^24^. Yet for most of these variants, their functional relevance remains unknown, mainly because experimental testing is resource-intensive and lagging far behind the pace of new variants are discovered, a common challenge in human genetics. For many disease-associated genes, most missense variants are currently classified as variants of uncertain significance (VUS), meaning that VUS substantially outnumber variants with confirmed pathogenic or benign effects^25^. Computational approaches to predict variant effects have become an important tool to address this gap.

In recent years, computational variant effect predictors have dramatically improved, especially with the advent of AI and evolutionary modeling. Deep generative sequence models, such as EVE, infer damaging variant effects from evolutionary conservation patterns without requiring labelled training data^26^. At the same time, large-scale AI-driven approaches such as AlphaMissense (a derivative of AlphaFold-based modeling) leverage protein language models and structure-based inference to predict functional effects of all possible human missense variants^27^. Many other predictors based on different principles are widely used to study variant effects. For example, GEMME^28^ estimates the impact of amino acid substitutions by quantifying evolutionary distance from the closest tolerated sequence neighbors. REVEL^29^ is a meta-predictor that combines features from multiple conservation-, structure-, and pathogenicity-oriented scores to yield a consensus classifier. DeMaSk^30^ has been trained on experimental deep mutational scan data and incorporates biophysical descriptors to identify substitutions that disrupt local packing or functional sites. Although these tools achieve high accuracy in classifying variants as neutral or deleterious, they typically yield a single score or classification, providing limited insight into the underlying molecular mechanism of dysfunction. In other words, a variant might be flagged as “likely damaging”, but it remains unclear whether this is due to protein misfolding, loss of an interaction interface, altered enzymatic activity, or some other structural perturbation^31^. To develop targeted hypotheses and guide follow-up experiments, it is valuable to pinpoint the mechanistic basis of a variant’s effect.

To address this limitation, structure-based mechanistic frameworks analyze variant effects across various biophysical and functional features of a protein. One such approach is MAVISp (Multi-layered Assessment of VarIants by Structure). This modular computational pipeline integrates diverse structural and sequence-based metrics to evaluate variant effects across multiple “mechanistic indicators”^32^. Rather than producing a binary classification, MAVISp evaluates how a variant might affect protein stability, local and long-range interactions, oligomerization interfaces, and phosphorylation sites. By considering the protein’s three-dimensional structure and dynamics, this approach can highlight *why* a variant might be deleterious, for instance, by destabilizing a critical helix or preventing a necessary protein–protein interaction. Such mechanistic predictions are particularly informative for large, multimeric proteins, such as ATG9A, whose function depends on coordinated lipid transport and protein complex formation.

In the present study, we applied the MAVISp structure-based variant assessment to human ATG9A to systematically characterize the effects of missense mutations and identify variants that could compromise ATG9A function. Using an *in silico* saturation mutagenesis approach, we evaluated every possible amino acid substitution at each position in multiple functional contexts: (i) the monomeric protein (to assess intrinsic folding stability and long-range allosteric effects), (ii) the homotrimeric assembly of ATG9A (to assess impacts on oligomerization interfaces and overall complex stability), and (iii) the ATG9A–ATG2A complex (to evaluate variants at the binding interface that might disrupt ATG2A-mediated lipid transfer). For each variant, we examined its predicted effects on protein stability (folding free energy), intra-protein interaction networks or allosteric communication paths, oligomer formation, and known regulatory sites (such as phosphorylation or ubiquitination sites on ATG9A). We further integrated predictions from four independent Variant Effect Predictors (VEPs; AlphaMissense, EVE, GEMME, and DeMaSk which provide an orthogonal assessment of how likely each variant is to be damaging^27,30,33,34^. Furthermore, we cross-referenced all variants with disease and cancer variant databases (ClinVar for clinically observed variants, COSMIC for somatic cancer mutations, and cBioPortal for tumor mutation frequencies in patient cohorts) to highlight variants already observed in biomedical contexts. This multi-layered approach enabled prioritizing candidate ATG9A variants predicted to have strong deleterious effects (e.g., loss-of-function or disruption of autophagy-related mechanisms) for future experimental validation.

## Results and Discussion

### Overview of the data used in the study

We carried out *in silico* saturation mutagenesis and retrieved annotations for disease-related variants from ClinVar^35^, COSMIC^22^, and CbioPortal^36^. The initial three-dimensional (3D) structures used to analyze the protein in its free form, in its multimeric state, and in complex with ATG2A are shown in **Figure 1**. The monomeric structure was used to assess structural stability, long-range interactions, and the relationship between variant effects and phosphorylation sites. The dimeric interfaces forming the trimeric assembly of ATG9A were used to estimate the impact of variants on interactions relevant to multimerization. Finally, the structure of the ATG9A-ATG2A complex was analyzed to predict variant effects at the binding interface. All analyses were carried out in both the *simple* and *ensemble modes* of MAVISp across all mechanistic indicators, except for local interaction assessments of ATG9A and ATG2A in *ensemble mode*, which were excluded due to computational limitations (**Figure 2)**.

**Figure 1.**
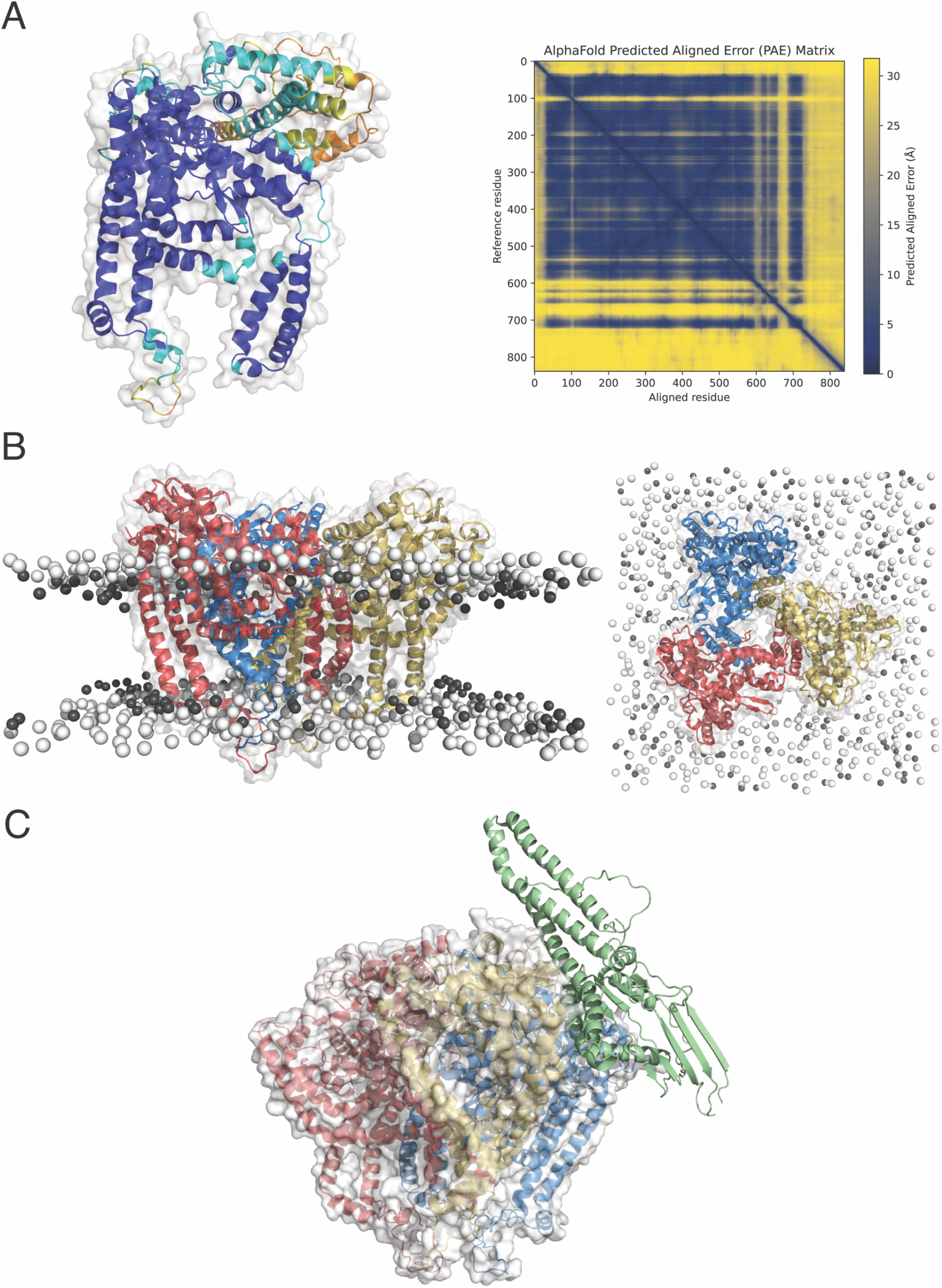
Overview of the structures used for the MAVISp assessment. (**A**) AlphaFold model of human ATG9A (UniProt: Q7Z3C6) in its free state, used throughout the MAVISp assessement to examine structural stability, long-range interactions, and the interplay between variant effects and phosphorylation sites.The model is color-coded by confidence: dark blue indicates very high confidence (pLDDT >90), light blue high confidence (90 > pLDDT >70), yellow low confidence (70 > pLDDT >50), and orange very low confidence (pLDDT <50). The panel on the right shows the Predicted Aligned Error (PAE) matrix, which represents the expected positional error (in Å) between residues. Lower values (dark blue) correspond to higher confidence in relative positioning, whereas higher values (yellow) indicate greater uncertainty. The axes correspond to the residue indices of ATG9A. The structure displayed corresponds to the truncated form used in our analysis (ATG9A_36–522_), while the PAE matrix refers to the full-length construct (ATG9A_1-839_). (**B**) Trimeric model of ATG9A embedded in a POPC:CHL (3:1) lipid bilayer. Each ATG9A_36–522_ monomer is represented in a distinct color (promoter A in red, B in blue, and C in yellow). Lipids are shown as spheres. The right panel provides a top view of the trimeric assembly. These structural models were used to analyze the dimeric interfaces that stabilize the trimeric assembly of ATG9A and to estimate the impact of sequence variants on interactions relevant to multimerization.(**C**) Model of the ATG9A–ATG2A complex (ATG9A_34–723_ and ATG2A_1512–1934_) from vanVliet & Chiduza et al. (2022). The three ATG9A protomers are colored as in panel B (A, red; B, blue; C, yellow), and ATG2A is shown in green. This structure was used to predict the effects of variants at the ATG9A–ATG2A binding interface.

**Figure 2.**
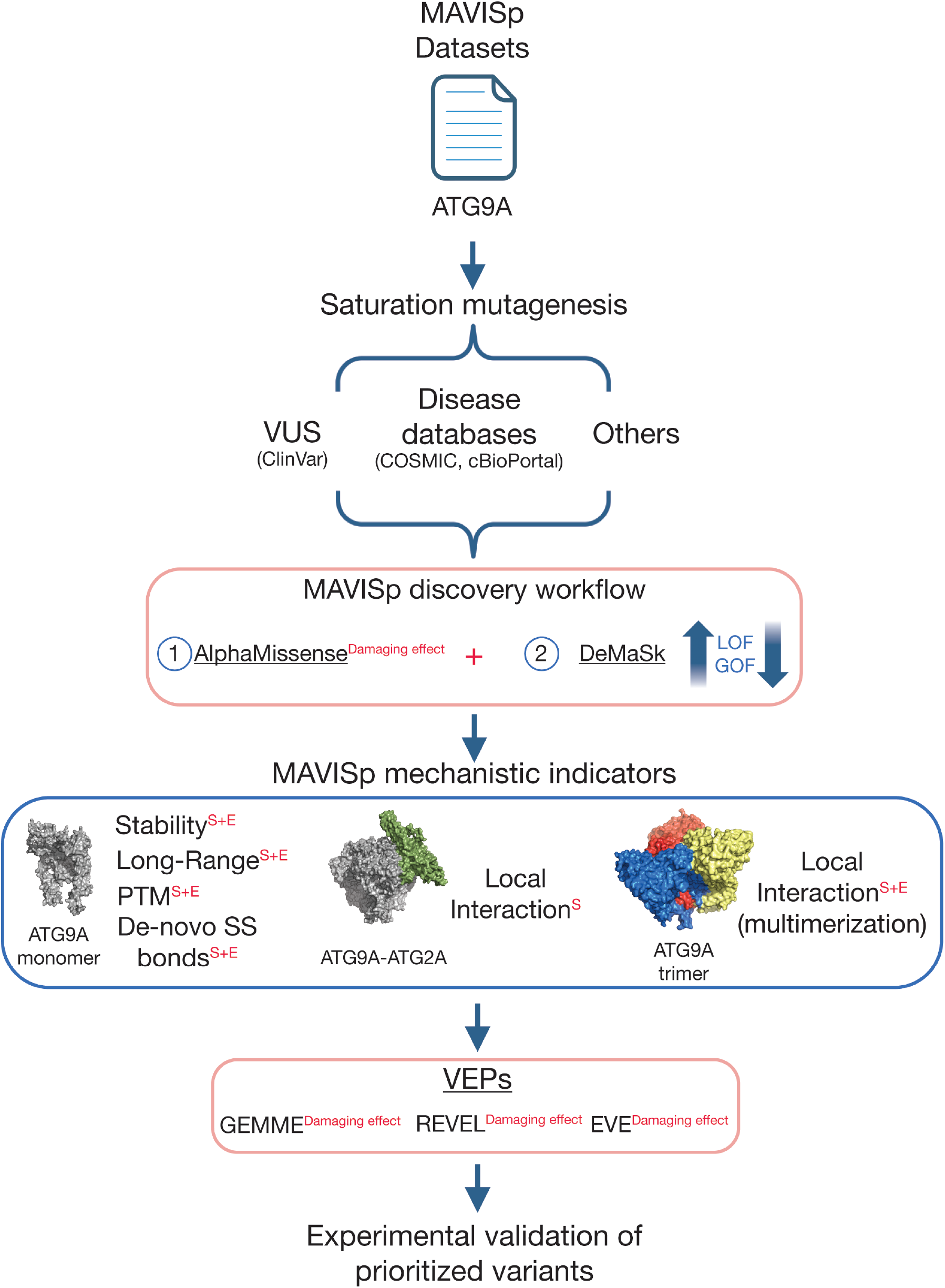
Overview of the workflow applied in this study to identify damaging ATG9A variants for experimental validation. At first, we collected data using most of the modules available in the MAVISp framework for ATG9A in both *simple* (S) *and ensemble modes* (E), applying saturation mutagenesis to investigate all missense variants with structural coverage. The framework also allowed the inclusion of annotations for variants reported in three disease-related databases, i.e., ClinVar, COSMIC, and cBioPortal. For ATG9A, only one benign variant is reported in ClinVar, and the remaining variants we studied are classified as Variants of Uncertain Significance (VUS) or are variants not yet reported in ClinVar. We thus apply the default MAVISp discovery workflow to identify candidates for experimental validation based on the usage of Variant Effect Predictors (VEPs). This included a first step in which variants reported as damaging by AlphaMissense were retained, followed by a step in which we applied DeMaSk. In the second step, DeMasK is used to maintain variants with a possible loss- or gain-of-fitness phenotype. For variants that pass VEP-based filtering, we used the MAVISp structure-based module to identify potenEal mechanisEc indicators. We used the structure of the ATG9A monomer for stability, long-range effects, and interplay between phosphorylation (as post-translational modification, PTM) and amino acid substitutions in *ensemble mode*. The same monomeric structure is used in the *simple mode* for the evaluation of the alteration of native or de novo formation of disulfide bridges. Finally, each dimeric interface within the ATG9A trimer has been used to assess changes in local interactions in *ensemble mode*, whereas the ATG9A–ATG2A complex has been used in *simple mode*. Finally, the predictions from the other VEPs implemented in MAVISp (i.e., GEMME, REVEL, and EVE) are also verified on the selected candidates.

In accordance with the MAVISp protocol, we also included predictions of damaging effects from four different VEPs, i.e., AlphaMissense^27^, EVE^33^, GEMME^28^, and REVEL^29^, which may aid in identifying potentially pathogenic variants. Furthermore, we applied DeMaSk^30^, which provides predictions of the loss/gain in fitness upon amino acid substitution (**Materials and Methods**). MAVISp also allows the integration of experimental classifications or scores for each mechanistic indicator. To this end, we curated literature-based experimental data (see Materials and Methods) and identified the 15 variants listed in **Table 1**.

**Table 1.**
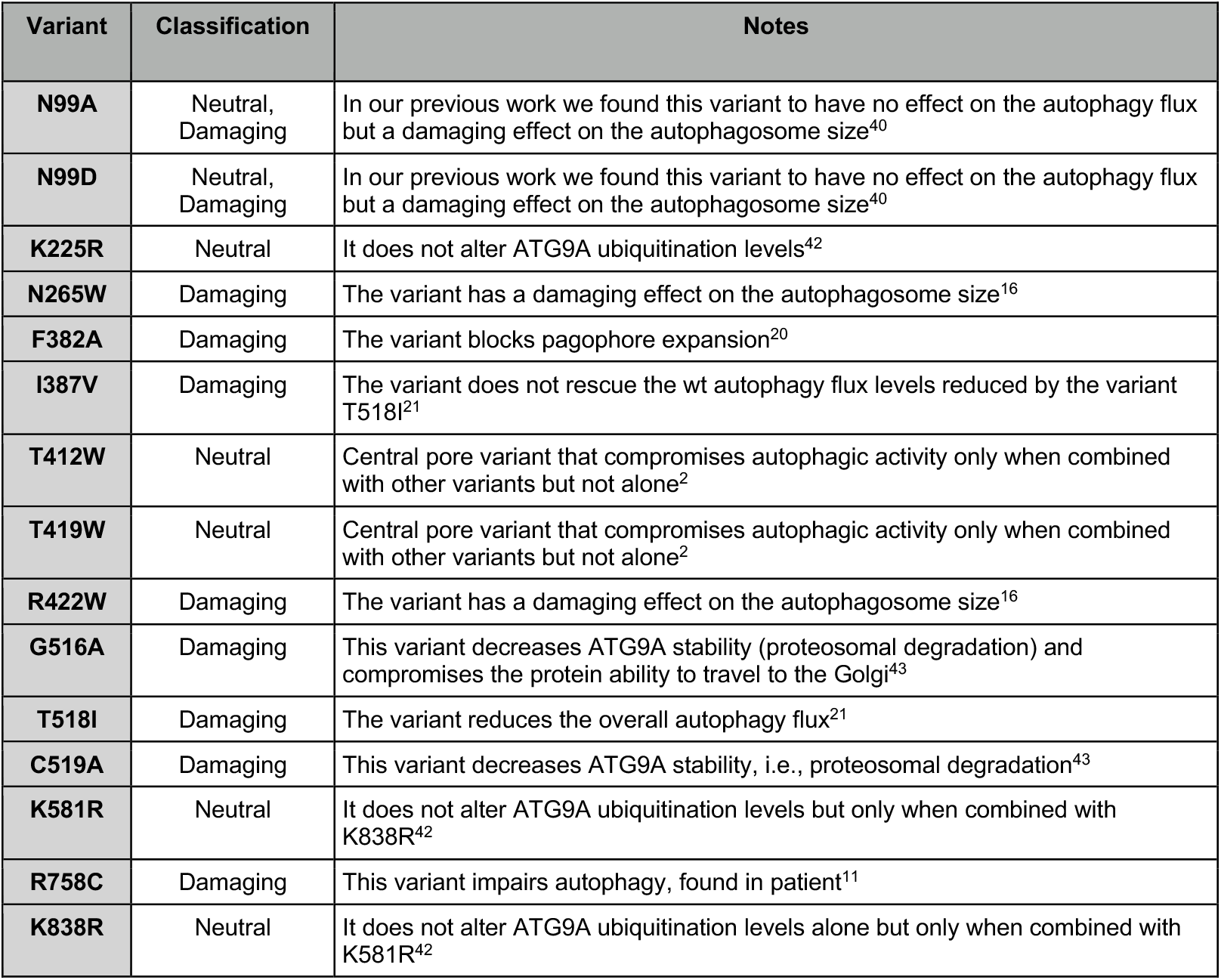
Experimentally validated single variants of ATG9A have been reported in the literature. The first column reports the specific variant, the second column indicates its functional classification (Neutral or Damaging) based on the experimental results, and the third column provides additional notes on the observed biological effects and the corresponding reference to the original study in which the experiments were performed.

Unless otherwise stated, the results discussed in the text refer to *ensemble-mode predictions* for phosphorylation, long-range interactions, multimerization, and stability. For the ATG9A-ATG2A interaction, the results are based on the *simple mode* assessment. The advantage of using the *ensemble mode* in this case study is that the protein remains embedded in the membrane during molecular dynamics simulations, providing a more realistic environment for evaluating variant effects.

### ATG9A damaging variants and associated molecular mechanisms

We used the MAVISp framework to predict variant effects, identify potentially damaging variants, and associate them with mechanistic indicators, thereby prioritizing candidates for experimental validation.

Among the ClinVar entries, only one variant (i.e., A663V) had structural coverage and a review status of 1. The aggregated MAVISp results confirmed that this variant has neutral effects (**Figure S1**).Among the variants reported in COSMIC (i.e., 156 and 80 variants in simple and *ensemble mode*, respectively) and cBioPortal (i.e., 194 and 113 variants in *simple* and *ensemble mode*), we identified one variant with a damaging effect, loss of function, and altering structural stability, i.e., G357D (**Figure 3A-C**). G357D is reported in both cancer databases. In particular, it has been identified in the colorectal cancer cell line LoVo, derived from a metastatic adenocarcinoma and frequently used as a model for colorectal cancer^37,38^. According to UniProt, residue G357 lies within a cytoplasmic topological domain of ATG9A. It is located near a known ubiquitination site (K359), suggesting that the mutation could influence post-translational regulation. The variant was not reported in previous experimental studies on its effect on ATG9A function; therefore, it was selected for further investigation in this work. Similarly, we selected L168P (**Figure 3A-C**), reported in ClinVar as a VUS with a review status of 1. No specific pathological condition has been associated with this variant; the residue is positioned within the cytoplasmic region of the protein and in proximity to a transmembrane helix, suggesting that the mutation could affect local folding and was therefore selected for further investigation in this work. In the disease-related databases used by the MAVISp protocol, we also identified R211C and R211H as variants with potentially damaging and loss-of-fitness effects located in regions important for early folding events (**Figure 3A-C**).

**Figure 3.**
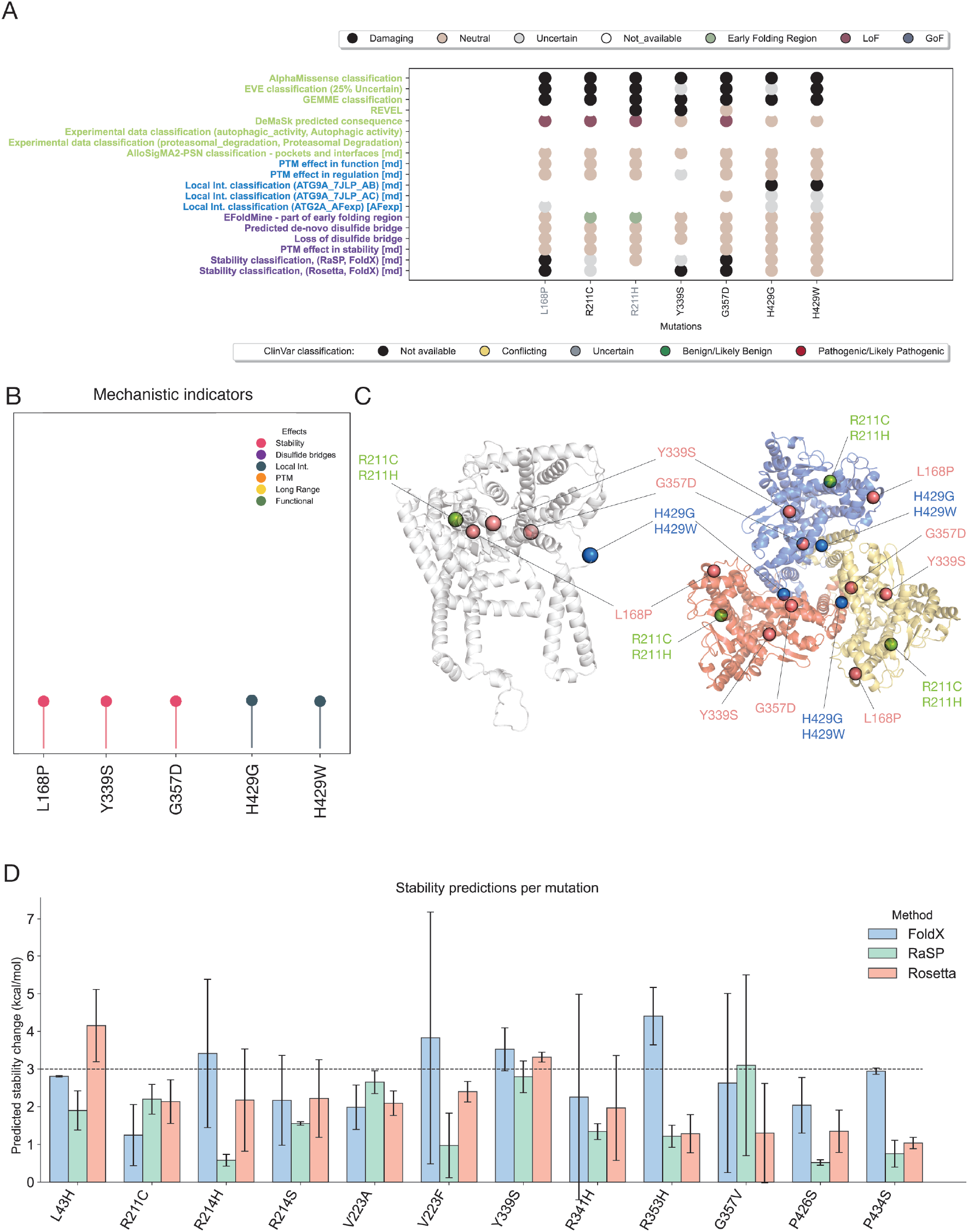
MAVISp predictions of damaging ATG9A variants and associated mechanistic indicators. Dot plot summarizing the MAVISp predictions for the seven variants identified as destabilizing for at least one mechanistic indicator (L168P, R211C, R211H, Y339S, G357D, H429G, and H429W).Lolliplot representation of the five variants for which mechanistic indicators were assigned (excluding R211C and R211H, which were associated with early folding events). **(C)** Structural mapping of the seven variants on ATG9A. Leb: monomeric structure of ATG9A (residues 36–522, cartoon, white). Right: trimeric assembly (residues 36-522, cartoon; chain A in red, chain B in blue, chain C in yellow). Variants are shown as spheres and colored according to their predicted mechanistic effect: red, altered structural stability; blue, perturbed local interactions; green, early folding events. **(D)** Grouped bar plot showing the predicted folding ΔΔG values of the 12 variants reported in disease-related databases that were classified as uncertain for stability by the MAVISp consensus approach (FoldX + RaSP). Bars represent predictions from FoldX (blue), RaSP (green), and Roseha (red), with error bars indicating the standard deviation across ensemble calculations. All ΔΔG values are reported in kcal/mol. The dashed horizontal line at 3 kcal/mol denotes the threshold used to classify variants as destabilizing.

Furthermore, we identified 12 variants reported in disease-related databases that, according to the MAVISp consensus approach, exhibited uncertain stability predictions and showed a signature of damaging effects for AlphaMissense (**Table S1**). The average folding free energy values used for classification were overall representative of the estimated impact of each conformation in the ensemble used for the analyses, as evidenced by low standard deviations (**Figure 3D, Table S1**), except for G357V. Of these, we noticed that R214H, V223F, Y339S, and R353H were predicted to be damaging by FoldX but not by RaSP (**Figure 3D, Table S1**). We thus performed calculations using the Rosetta protocol implemented in MAVISp in *ensemble mode* to further verify these predictions, with Y339S resulting in destabilization (**Figure 3D, Table S1**).

Finally, we identified an additional 451 variants that were not yet reported in the disease-related resources used in this study, with predicted damaging effects, loss-of-fitness signature, and associated mechanisms (**Figures S2**). The variants were all expected to alter structural stability, except for A221C, which was predicted to promote de novo disulfide bridge formation with C519 (**Figure 4A**) and Y209D/P (**Figure 4B**). We also verified the distance between the functional groups of the residues at positions 221 and 519 in the three MD replicates of the wild-type ATG9A form to further consolidate the potential of disulfide bridge formation upon mutation predicted in the *simple mode*. Interestingly, in two of the three replicates, the two residues exhibit a distance distribution centered at 4 Å (**Figure 5A**).

**Figure 4.**
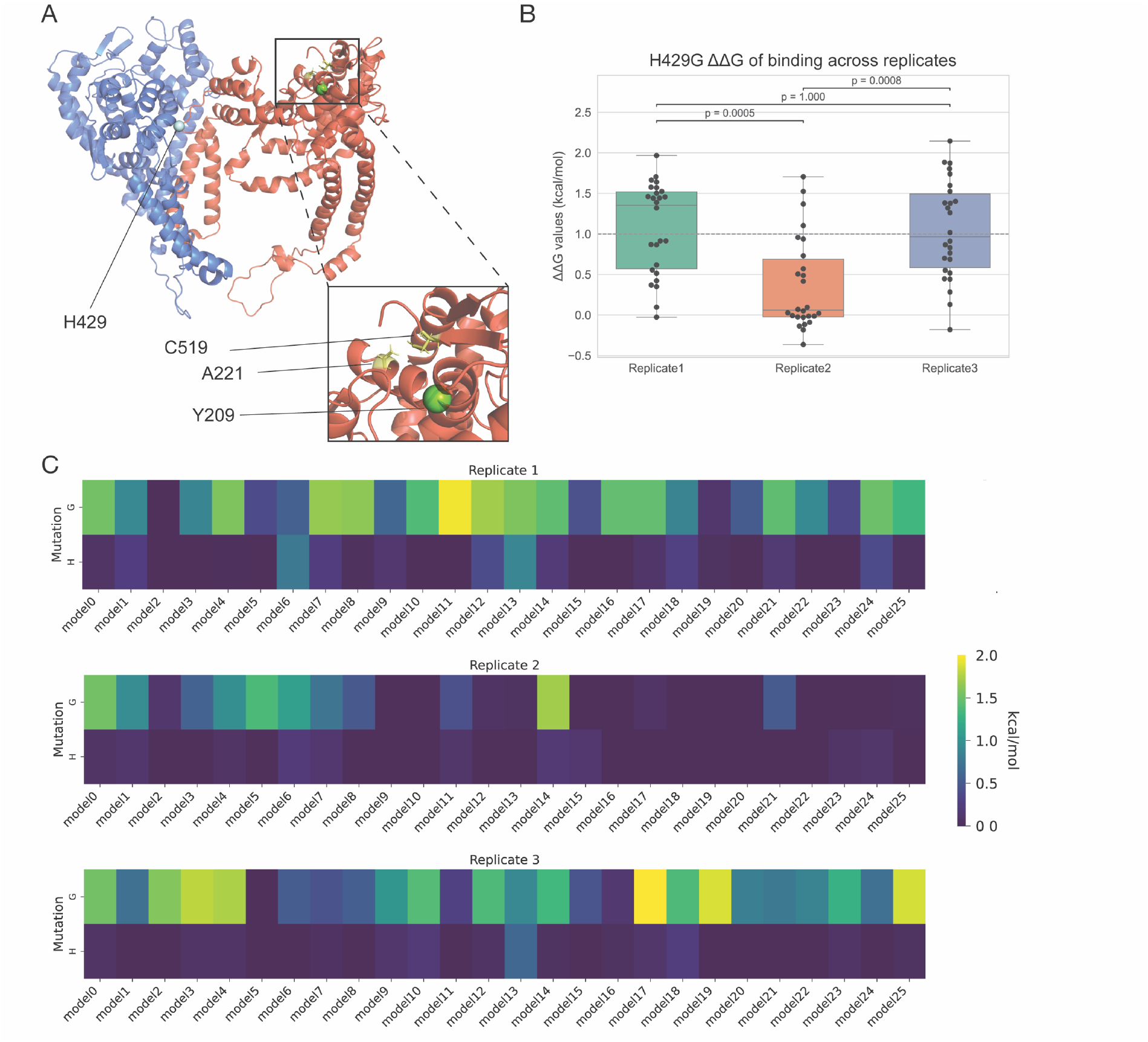
Structural mapping and MD-based evaluation of selected MAVISp-predicted ATG9A variants. (**A**) Mapping of the five selected variants on the AB interface of the ATG9A trimer used for the MD simulations. H429G and H429W (cyan sphere) are predicted to alter the subunit–subunit interface and potentially destabilize multimeric assembly. Y209D and Y209P (green sphere) are predicted to destabilize the local structure relative to the phosphorylated state of this conserved tyrosine. A221C (yellow sEcks) is predicted to promote a de novo disulfide bridge with C519 (also shown as yellow sEcks). The trimer is shown in cartoon representation (chain A in red, chain B in blue). Insets provide zoomed views of the environments surrounding Y209 and A221. (**B**) Box- and-swarm plot of the predicted binding ΔΔG values (kcal/mol) for H429G, computed with FoldX on structural ensembles extracted from the three MD replicates. Green, orange, and blue correspond to replicates 1, 2, and 3, respectively. Pairwise Mann–Whitney U tests were performed, and Bonferroni-corrected p-values are shown. (**C**) Heat maps reporting per-model binding ΔΔG values (kcal/mol) for H429G and the self-mutation H429H across all MD replicates. Each column represents an individual model from a given replicate, highlighting replicate-specific variability and the neutral behavior of H429G observed in replicate 2.

**Figure 5.**
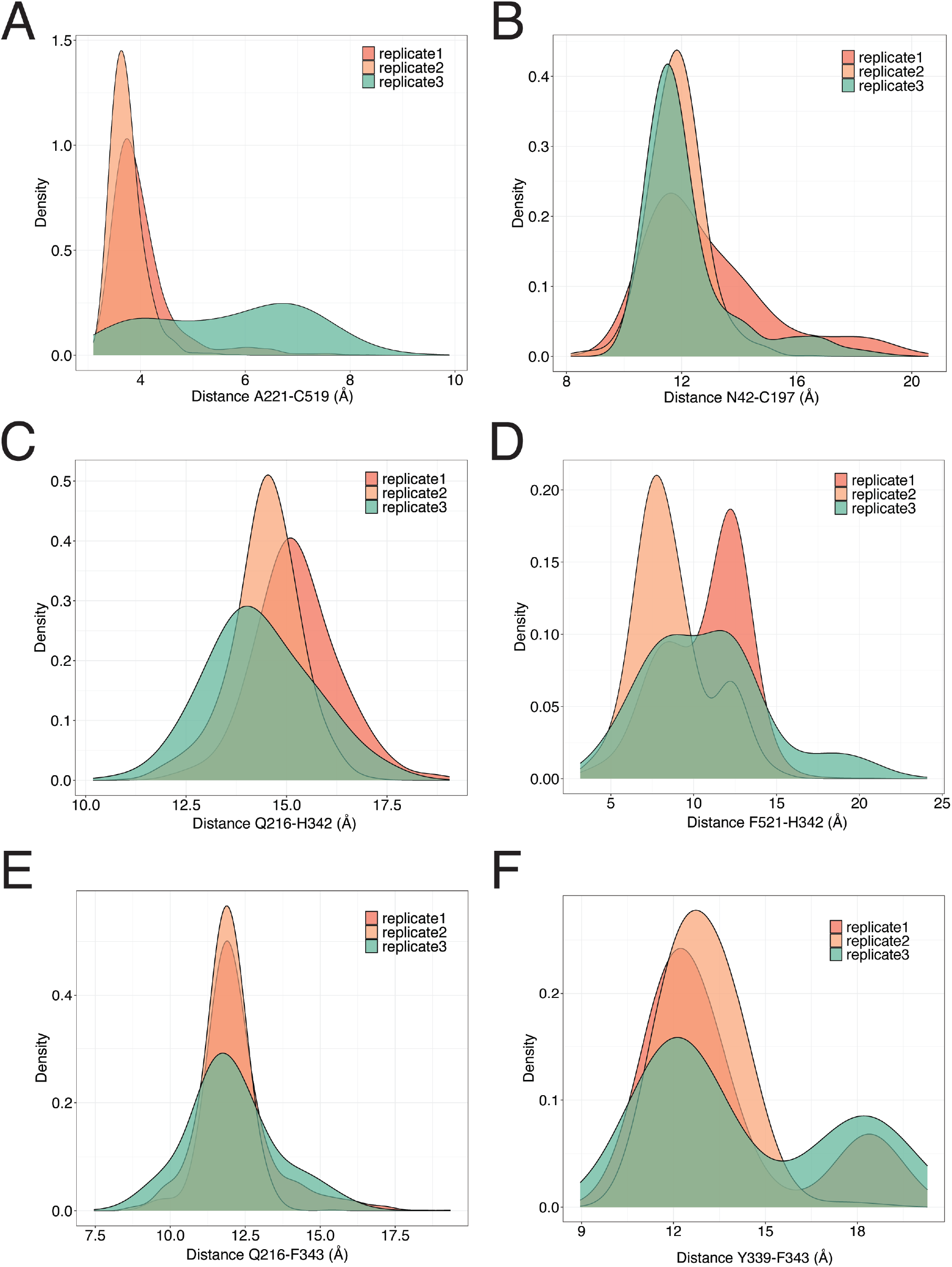
Distributions of inter-residue distances across MD replicates. The figure shows density plots of the distributions of selected pairwise distances between specific functional groups for each residue pair across the three MD replicates, shown in red (replicate 1), orange (replicate 2), and green (replicate 3). (A) Distance between the CB atom of A221 and the SG atom of C519. (B) Distance between the OD1 atom of N42 and the SG atom of C197. (C) Distance between the OE1 atom of Q216 and the NE2 atom of H342. (D) Distance between the CZ atom of F521 and the NE2 atom of H342. (E) Distance between the OE1 atom of Q216 and the CZ atom of F343. (F) Distance between the OH atom of Y339 and the CZ atom of F343.

Y209D/P are expected to alter a conserved tyrosine residue in ATG9A and to destabilize the structure relative to the phosphorylated state at this site (**Supplementary Movies 1-3**). To our knowledge, this site has not been detected in human phospho-proteomics datasets, and no cellular studies have investigated its potential regulation. Although large-scale phospho-proteomics studies report multiple phosphorylated peptides within ATG9A (PRIDE/ProteomeXchange datasets PXD017871, PXD053554), none include the segment containing Y209. However, Y209 is evolutionarily conserved in mammals, and a mass-spectrometry study in mouse atg9a has reported phosphorylation at the equivalent residue^39^ suggesting that this position may be phosphorylated under specific conditions in vivo. In support of this idea, it is predicted to have a score >0.8 as a possible phosphosite (see Materials and Methods). Nevertheless, we observed that the site is located in a small α-helix that points towards a buried region (**Supplementary Movies S1-3**) and exhibits low solvent accessibility during the MD simulation (average side-chain solvent accessibility = 19.8%). A longer timescale simulation might be required to probe whether this is a cryptic phospho-site that becomes accessible to kinases through conformational changes in the disordered regions around it.

We also identified 59 variants at six different sites (C197, F343, F521, H342, M448, and N344) with predicted long-range effects to response sites in protein pockets. Nevertheless, the majority of these pairs of targets and response sites are in regions that, upon local conformational changes, would bring the side chain of the mutation and the response site into direct contact and are thus unlikely to underlie a relevant allosteric effect. To verify this hypothesis, we calculated the distances between the functional groups of mutation sites and target sites in the three MD replicates of the ATG9A wild-type form (**Figure 5B-F**). Among the six positions highlighted by the LONG_RANGE module, only C197, H342, and F343 were analyzed in this way, as they were the only sites *not* already in direct contact with their corresponding response sites in the structure used for the initial assessment. The remaining sites (M448, N344, and F521) lie adjacent to their predicted response residues in the reference structure, making a long-range perturbation structurally implausible. For all three sites, the analysis confirms that the distributions are centered well above 10 Å (**Figure 5B-F**), indicating that neither stable nor transient interactions occur during the trajectories. The only partial exception is the H342– F521 pair (**Figure 5D**), where the center of the distribution varies across replicates, approximately 8 Å in replicate 1, ∼13 Å in replicate 2, and a broader distribution spanning 8–13 Å in replicate 3. Although this pair comes into closer contact in specific frames of the MD trajectory, the distances remain beyond the range required for short-range interaction (<5–6 Å).

By relaxing the criteria for identifying mechanistic indicators without using the second layer of information provided by DeMaSk (**Figure S3**), we identified H429G and H429W, which can potentially affect the interaction between ATG9A subunits at the interface **(Figures 3A-C**) and destabilize multimeric assembly.

We also verified the predictions using two additional ATG9A MD replicates for the subset of variants selected for experimental validation. We observed good overall agreement among MD replicates for variants predicted to affect structural stability (**Table S2**). On the contrary, in the case of H429G, the conformations explored during replicate 2 suggest a neutral effect for this variant at the interface (**Figure 4A-C**). This is in line with our previous MD results on ATG9A^40^, where different replicates sampled distinct open/closed interface configurations, and some trajectories transiently stabilized closed-like states.

To better understand why the predicted effect of H429G differs across different MD replicates, we considered the binding ΔΔG upon the glycine substitution at position 429 for each of the 25 models extracted from each MD simulation and used for FoldX calculations in the MAVISp *ensemble mode*. The results are reported in the boxplot and heatmaps in **Figure 4B-C**. Additionally, we estimated the number of initial conformations for which a destabilizing binding ΔΔG was obtained. We observed that MD replicates 1 and 3 have a predicted damaging effect in approximately 54% of models, whereas replicate 2 has a predicted damaging effect in only 15% of cases. This difference is consistent with replicate 2 sampling a more closed interface, in which the local environment around H429 is more constrained, and the contacts remain less perturbed by the mutations (**Figure 4B-C**). These results indicate a conformation-dependent role impact for the amino acidic substitutions at the H429 site at the monomer-monomer interface of the trimeric complex of ATG9A.

Considering the above analysis, we prioritize L168P and G357D for future experimental validation, given their potential impact on structural stability and their reported status as VUS or in cancer databases, respectively. Additionally, we included the variant H429G given its conformation-dependent effect on local interactions at the dimerization interface of the ATG9A multimeric assembly. Finally, we compared the results from the different VEPs implemented in MAVISp for each of the variants selected for experimental validation (**Figure 6**). L168P was consistently classified as highly damaging across all VEPs. This is in line with the expected structural penalty associated with introducing a proline within a helical region. G357D is classified as potentially damaging by most VEPs, suggesting that introducing a charged residue at this position is poorly tolerated.

**Figure 6.**
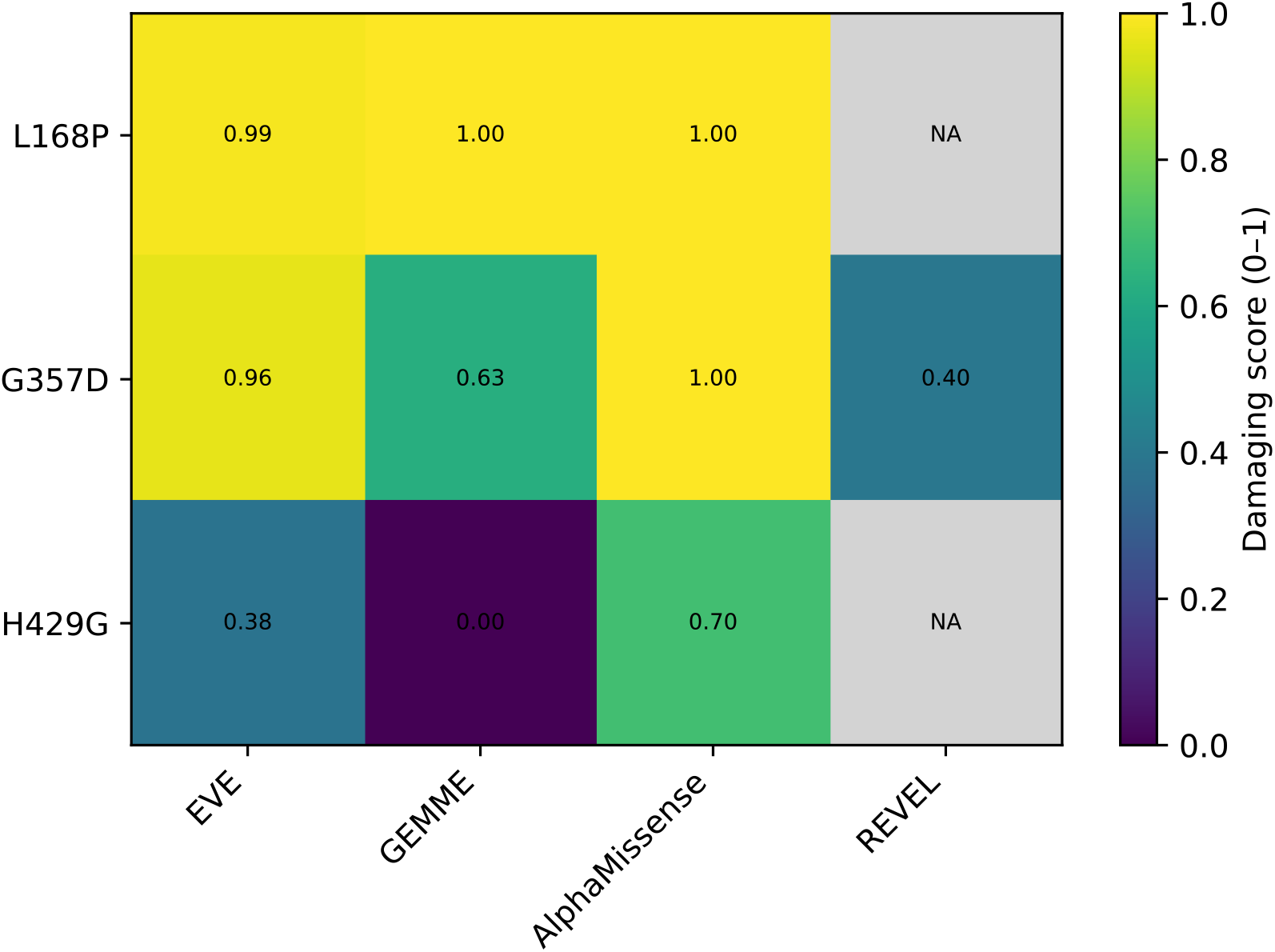
Variant effect predictions for selected ATG9A mutants. The heatmap summarizes the results from the four variant effect predictors (VEPs) implemented in MAVISp for the three ATG9A variants chosen for future experimental validation (i.e., L168P, G357D, and H429G). The panel reports the scores for EVE, AlphaMissense, REVEL, and normalized GEMME scores (see Materials and Methods). Missing REVEL data are indicated in grey.

In contrast, H429G is predicted to have a damaging effect only by AlphaMissense and a lower impact by GEMME and EVE. The structure-based analyses conducted with MAVISp suggest a conformation-dependent impact for this variant, which may perturb the monomer–monomer interface and thereby affect multimerization. Because this variant represents a case where the predicted VEPs do not well capture structural effects, we would need to evaluate the variant experimentally in the future, despite the variant not yet being identified in disease-related resources.

## Conclusions

In this study, we systematically analyzed the functional landscape of missense variation in the human autophagy regulator ATG9A by integrating in silico saturation mutagenesis, Variant Effect Predictors, and a structure/dynamics-based mechanistic assessment of variant effects. By applying the MAVISp framework across multiple structural states of ATG9A, the monomer, the homotrimer, and the ATG9A–ATG2A complex, we were able to dissect how individual amino acid substitutions might perturb structural stability, local contacts, long-range communication, and oligomerization of ATG9A. This multi-layered analysis revealed different potentially damaging variants, many of which had not been previously documented in disease-related databases.

Our work highlights several mechanistic classes through which ATG9A dysfunction may arise, including destabilization of the protein structure, altered trimeric assembly, and perturbation of potential regulatory sites, such as phosphorylation. Importantly, integrating these structure-based mechanistic indicators with evolutionary and deep-learning Variant Effect Predictors enabled us to prioritize high-confidence variants predicted to compromise ATG9A function. These findings underscore the value of mechanistic prediction frameworks for narrowing the gap between variant discovery and functional interpretation.

More broadly, our study provides the first comprehensive structure-informed atlas of missense variation in ATG9A. This resource will aid researchers in evaluating ATG9A variants encountered in population sequencing, cancer genomics, and rare disease diagnostics. From a basic research perspective, these data provide a catalog of variants to probe how specific residues, their structural elements, and interface regions contribute to ATG9A’s lipid-scrambling activity, membrane-remodeling properties, and assembly into higher-order complexes. Although ATG9A’s lipid-scramblase activity has been demonstrated by structural and biochemical studies^15,17^, pinpointing residues that specifically govern scrambling, rather than contributing more broadly to protein stability or autophagy function, remains challenging. Most experimentally tested variants disrupt autophagy or protein stability overall, making it difficult to isolate effects on lipid translocation itself^15^. Our analyses provide a selection method for variants neutral to stability to study for functional readouts. Of note, this work contributed to the inclusion of a new module for MAVISp to assess the effects of variants on the disruption or de novo formation of disulfide bridges, which can be extended to all protein entries in the database. Finally, we observed that the current design for the long-range module in ensemble mode could be improved by introducing a step to verify the distance distribution between the target and response sites, to avoid assigning long-range damaging effects to residues that may come into contact through local conformational changes. The approach introduced here is generalizable and can be readily applied to other autophagy factors or membrane proteins whose dysfunction contributes to human disease. By bridging high-throughput computational predictions with mechanistic insight, this work establishes a foundation for elucidating the molecular consequences of genetic variation in autophagy and for prioritizing variants for deeper investigations.

## Methods

### Identification and aggregation of variants for the study

ATG9A variants were collected from the disease databases supported by MAVISp, through Cancermuts^41^ (https://github.com/ELELAB/cancermuts), which provides an automated framework for querying, harmonizing, and annotating mutational data. Variants were retrieved from COSMIC^22^ and cBioPortal^36^ to identify cancer-related variants, and from ClinVar^35^. We used the RefSeq NP_001070666 for the search in ClinVar, corresponding to the main UniProt isoform, verifying the match with the uniprot2refseq tool of MAVISP (https://github.com/ELELAB/mavisp_accessory_tools/tree/main/tools). We used a dedicated Python script (*clinvar*.*py*, available in the mavisp_templates directory of https://github.com/ELELAB/MAVISp_automatization) to query the ClinVar data, which are then provided to Cancermuts for aggregation with COSMIC, cBioPortal, and other annotations at the sequence level. The data used for the final version of this publication refers to the updated collection performed on the 17th of September 2025 to ensure up-to-date coverage of variant annotations. Variant retrieval and annotation were performed using the Cancermuts workflow.

### Annotations from experiments

We included curated annotations from published functional studies in the EXPERIMENTAL DATA module of MAVISp, reporting whether each ATG9A variant exhibited damaging or neutral effects on proteasomal degradation or autophagic activity. The literature search identified experimentally characterized single–amino acid variants, including N99A and N99D, which were previously shown to be neutral for autophagy flux but damaging to autophagosome size^40^; K225R, K581R, and K838R, which do not alter ATG9A ubiquitination levels^42^; N265W and R422W, which impair autophagosome size^16^; F382A, which blocks phagophore expansion^20^; I387V and T518I, which reduce autophagy flux^21^; T412W and T419W, which affect autophagic activity only in combination with other pore variants^2^; G516A and C519A, which decrease ATG9A stability and trafficking^43^; and R758C, a patient-derived variant known to impair autophagy^11^. These experimentally validated classifications (damaging or neutral) are summarized in **Table 1** and incorporated into MAVISp for comparison with predictions.

### Variant Effect Predictors used in the study

We applied the default discovery workflow of MAVISp, since for this protein, there are no ClinVar data to calibrate VEPs, as we did in other focused studies^44,45^. The discovery workflow includes an assessment based on the AlphaMissense score^27^ to identify potentially damaging variants, and a second layer of investigation based on DeMaSk^30^ to further retain only variants with a signature of loss or gain of fitness. A more relaxed approach is to consider only the results from the first step, without the filtering based on DeMaSk.

We then analyzed the results from all available VEPs implemented in MAVISp for the subset of variants of interest for experimental validation, including REVEL^29^, GEMME^28^, and EVE^26^. The thresholds used for each VEP are discussed in detail in the original MAVISp publication^32^. We noted that for R211C, genomic coordinates were included in the MAVISp output file, but with the wrong genome build. The correct genome built is hg38, and the corresponding REVEL score is 0.693.

### Comparison among VEP scores

To enable the integration of diverse variant effect predictors, GEMME ΔΔE scores, in which negative values correspond to reduced fitness, were converted to a unified damaging scale. Scores were multiplied by –1 and min–max normalized using the empirical minimum and maximum values observed across the full ATG9A mutational dataset. The resulting 0–1 value reflects relative damaging potential, with 1 indicating the most deleterious prediction. EVE, AlphaMissense, and REVEL scores were used in their native 0–1 format.

### Structures used for analyses and simulations

We used the STRUCTURE_SELECTION module of MAVISp^32^ to retrieve the AlphaFold (AF) model of human ATG9A from the AlphaFold Protein Structure Database^46^, together with information on available experimental structures in the Protein Data Bank (PDB)^47^ through PDBMiner^48^. We aimed to reuse previously published ATG9A simulations for the analyses^40^ and to verify that the starting structure meets the criteria for inclusion in MAVISp entries. To do so, we used the tools within the STRUCTURE_SELECTION module of MAVISp, especially the quality control based on Procheck^49^, which resulted in only 0.2% of residues in the disallowed regions of the Ramachandran plot, confirming the overall quality of the initial structure used for simulations.

To model ATG9A for MAVISp analyses, we used both predicted and experimental structures. For the *ensemble mode*, the starting structure is the cryo-EM structure of the human ATG9A homotrimer in its closed conformation or the monomeric chain extracted from it, depending on the class of calculations (PDB 7JLP^2^). As described in our previous study on ATG9A glycosylation^40^, this model was refined by selecting residues 36–522, reconstructing missing coordinates within residues 96–108 using MODELLER, and evaluating the resulting models with SOAP-Protein-OD and DOPE-based metrics. In the present work, we used the same curated cryo-EM–based model, omitting glycosylation, as the current analysis focuses exclusively on the non-glycosylated protein. Additionally, we have also included data following the default MAVISp protocol in *simple mode*, which relies on the AlphaFold model of the monomeric human ATG9A (AF-Q7Z3C6-F1, downloaded on 05/09/2024) from the AlphaFold Protein Structure Database. The model was trimmed to remove regions with low pLDDT scores or high PAE values, resulting in a construct spanning residues 36–725 with high predicted confidence. We also collected data in the *simple mode* for the same experimental structure used in the simulations.

We used the three replicates of one microsecond each produced using all-atom classical Molecular Dynamics in explicit solvent with the CHARMM36m force field^50^ for ATG9A_36–522_ as starting conformational ensembles for the analyses and calculations with the LONG_RANGE, STABILITY, PTM, and LOCAL_INTERACTIONS modules from MAVISp in *ensemble mode*. Details of these simulations are provided in the reference^32^. We used both the AF model and the experimental structure for the analyses in the *simple mode* with the module DISULFIDE BRIDGES.

### Structural Analysis with MAVISp modules

The protocols for each of the MAVISp modules are extensively described in the original publication, except for the DISULFIDE BRIDGE module, which has been recently designed and applied for the first time in this study (see more details in the next Method Section). We used the STABILITY, LONG_RANGE, PTM, DISULPHIDE BRIDGES, LOCAL_INTERACTIONS, and LOCAL_INTERACTIONS_HOMODIMERS modules. Additionally, we included the annotations from the EFOLDMINE (early folding event sites) and SAS modules (solvent accessibility of side chains) based on analyses with EFoldMine^51^ and Naccess^52^, respectively. For these modules, we applied the scoring schemes and classification criteria established in the MAVISp framework. The STABILITY module reports changes in folding free energy (ΔΔG) computed with FoldX^53^, Rosetta^54^, or RaSP^55^. Variants are annotated as stabilizing (ΔΔG ≤ –3 kcal/mol), destabilizing (ΔΔG ≥ 3 kcal/mol), or neutral (–2 < ΔΔG < 2 kcal/mol), with substitutions in the intermediate ranges or showing method disagreement classified as uncertain. In *ensemble mode*, the folding free energy calculations were carried out using 26 frames extracted from the MD trajectories at regular times for MutateX^56^/FoldX^53^, or the structures of the three most populated structural clusters for RosettaDDGPrediction using the Cartesian protocol^54^ or its machine learning derivative, RaSP^55^. The clusters were obtained with GROMACS^57^ using the GROMOS clustering method, based on main-chain RMSD with a cutoff of 2.5 Å. The LOCAL_INTERACTIONS and LOCAL_INTERACTIONS_HOMODIMERS modules evaluate substitutions within 10 Å of an interaction interface, using FoldX and Rosetta to calculate changes in binding free energy and classify variants as stabilizing (ΔΔG ≤ –1 kcal/mol for both methods), destabilizing (ΔΔG ≥ 1 kcal/mol), or neutral (–1 < ΔΔG < 1 kcal/mol); cases lacking concordance or involving solvent-exposed residues with incomplete structural context are considered uncertain. The LONG_RANGE module employs the updated SBSMMA procedure implemented in AlloSigMA2^58^ to quantify allosteric responses. It assigns variants as stabilizing, destabilizing, mixed, or neutral, while substitutions associated with side-chain volume changes <5 Å^3^ are designated as uncertain. The PTM module integrates solvent accessibility, SLiM context, and predicted structural effects at phosphorylatable residues and follows the MAVISp decision logic to classify the variant impact. The DISULFIDE BRIDGE module identifies substitutions affecting native or predicted disulfide-bonding cysteines. EFoldMine annotations were incorporated using a threshold of 0.169 and requiring stretches of at least three consecutive early-folding residues, and solvent-accessibility values were derived from Naccess.

### Effect of variants on disulfide bridges and prediction of de novo disulfide bridges

We hereby introduce a new module within the MAVISp framework to evaluate the impact of the variants that alter a wild-type cysteine or introduce a cysteine upon mutation on the disruption and formation of disulfide bridges, respectively. This module performs two classes of analyses: i) identification of the pair of cysteines potentially involved in native disulfide bonds in the wild-type PDB structure used for the *simple mode*, and ii) prediction of potential *de novo* disulfide formation across models of the variants that introduce a cysteine upon mutation. The code to generate the data for this module is available in the GitHub repository https://github.com/ELELAB/mavisp_accessory_tools. For the first step, we analyzed the 3D structures used for the *simple mode* STABILITY module and five models of the same wild-type variant generated by FoldX^53^ using the MutateX protocol^56^ to predict pairs of cysteines which, based on geometrical criteria, could form a disulfide bond. All cysteine residues are extracted from the structure, and pairwise Cβ–Sγ geometries are evaluated according to established stereochemical criteria for covalent disulfides (refs). As a default, a cysteine pair is classified as a native disulfide bridge when the inter-sulfur distance (Sγ–Sγ) falls within 1.95–2.25 Å, and the corresponding bond angles (Cβ–Sγ–Sγʹ and Sγ–Sγʹ–Cβʹ) lie between 85° and 130°. An additional dihedral check (χ_SS ≈ ±120°) ensures correct torsional geometry. We also included a looser criterion in which only distances are assessed with some buffer, with a range of 1.8-3.6 Å, which is what we applied in this study. Pairs satisfying these conditions are annotated as *native disulfides*. The use of models generated with FoldX (five in this study) aims to create conformational diversity in the rotameric states accessible to the mutation site and its surroundings, and to avoid using a single static model of the protein of interest. The module also includes another option based on a rotamer scan limited to the mutation sites, starting from the initial structure and without dependence on FoldX. For cysteine pairs separated by ≤ 10 Å between sulfur atoms, the module rotates each Sγ atom about its Cα–Cβ axis using a small rotamer library (−60°, 60°, 180°), computes all possible inter-sulfur distances, and retains the minimum. These two procedures allow us to test if the geometric criteria are met under side-chain rotations. If any rotamer combination satisfies the looser 1.8–3.5 Å distance window, the pair is labeled as *rotamer-reachable*, indicating that a disulfide bond could form upon minor local rearrangements. Variants that substitute any of the cysteines participating in these native disulfide bonds are then considered disruptive for disulfide bridge formation and classified as damaging within MAVISp. The second step investigates whether the variant can induce *de novo* disulfide bonds in variant structures generated by FoldX using the MutateX protocol. For each variant, the module recursively scans all available PDB models generated by FoldX (e.g, five models per variant in this study), identifying the possible disulfide bridges involving the new cysteine and any surrounding cysteines. Each pair is tested using two possible filters: i) strict criteria, identical to those used for native bonds (Sγ–Sγ = 1.95–2.25 Å, correct angles and χ torsion), to detect fully formed disulfide linkages; ii) potential (loose) criteria, applied when the strict test fails with a relaxed Sγ–Sγ distance range of 2.25–3.5 Å. Pairs meeting this condition are considered *pre-bonded* or *near-bond* configurations that could form covalent linkages after minor relaxation. Each model is labeled *positive* if it contains at least one cysteine pair satisfying either condition. For every variant, the total fraction of positive models is computed across the ensemble. A majority-vote scheme then classifies the variant as *de novo disulfide-forming* (and therefore *potentially damaging*) if ≥ 50 % of its models are positive; otherwise, it is classified as *neutral*.

### Downstream analyses and visualization of MAVISp data

We used the scripts and protocols from the downstream analysis toolkit of MAVISp (https://github.com/ELELAB/MAVISp_downstream_analysis) to facilitate the interpretation of the results and the selection of variants for further investigation. This included dot plot and lolliplot, which help visualize the main results from the MAVISp features and the mechanistic indicator classes, respectively. Additionally, we used bar plots to compare the predicted changes in folding free energy across the different methods used in the STABILITY module.

### Post-processing of MD Trajectories with PLUMED Driver

We processed the MD trajectories using the open-source, community-developed PLUMED library^59^, version 2.3^60^. We defined the following pairs of distances: CB(A221)–SG(C519), OD1(N42)–SG(C197), OE1(Q216)–NE2(H342), CZ(F521)–NE2(H342), OE1(Q216)–CZ(F343), and OH(Y339)–CZ(F343). The distances were evaluated for each frame of the MD trajectory, and distribution plots were generated from the timeseries.

## Supporting information

Figure S1, Dotplot for the A663V variant

Figure S2, Lolliplot for the additional 451 variants that were not yet reported in the disease-related resources used in this study

Figure S3, Lolliplot for the analysis without the DeMaSk filtering

Supplemental Movie 1, Movie of the 1st replicate of the MD simulation

Supplemental Movie 2, Movie of the 2nd replicate of the MD simulation

Supplemental Movie 3, Movie of the 3rd replicate of the MD simulation

## Declaration of Generative AI and AI-Assisted Technologies in the Writing Process

During the preparation of this manuscript, the authors used OpenAI ChatGPT (version 5.2) for assistance with language editing and clarity. All content generated with this tool was carefully reviewed, revised, and validated by the authors, who take full responsibility for the accuracy, originality, and integrity of the final published work.

## Data availability

All data generated or analyzed in this study are available through the Open Science Framework at https://osf.io/e297x/, unless otherwise stated.

