## Supplementary figures and images for "Decoding ATG9A Variation: A Comprehensive Structural Investigation of All Missense Variants"

### Figure S1, Dotplot for the A663V variant

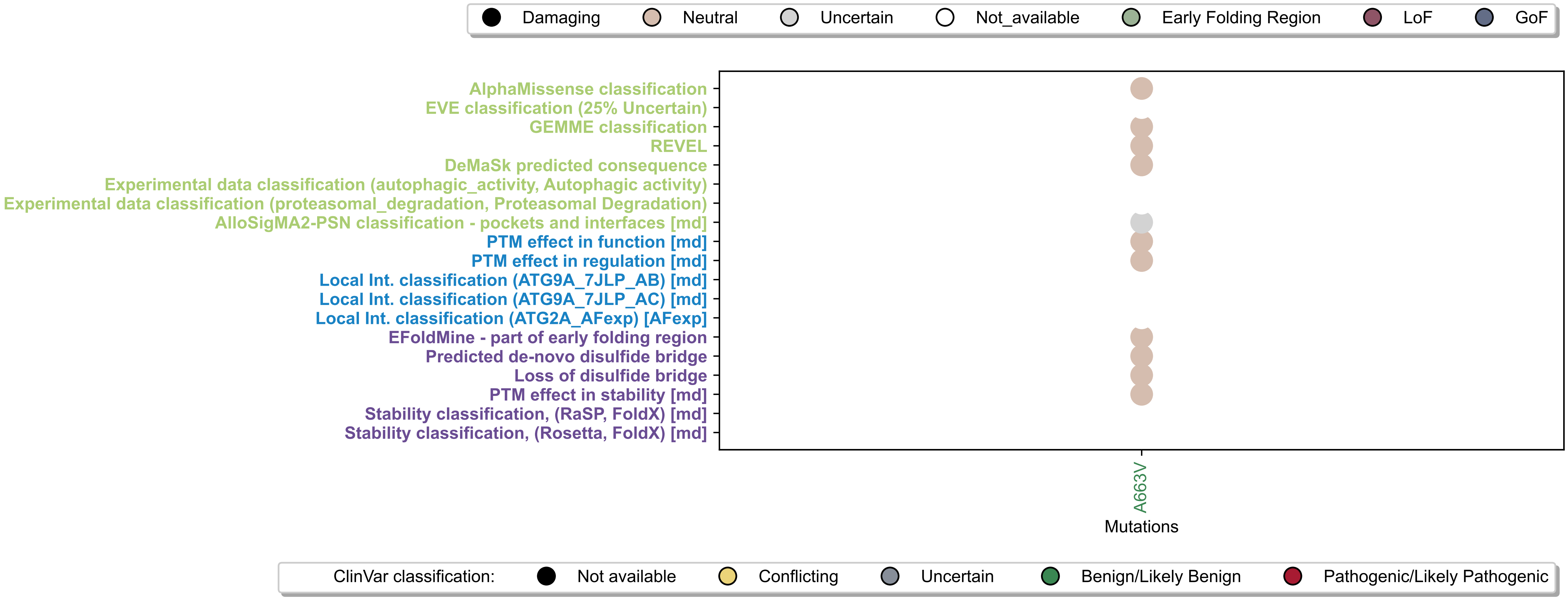

### Figure S2, Lolliplot for the additional 451 variants that were not yet reported in the disease-related resources used in this study

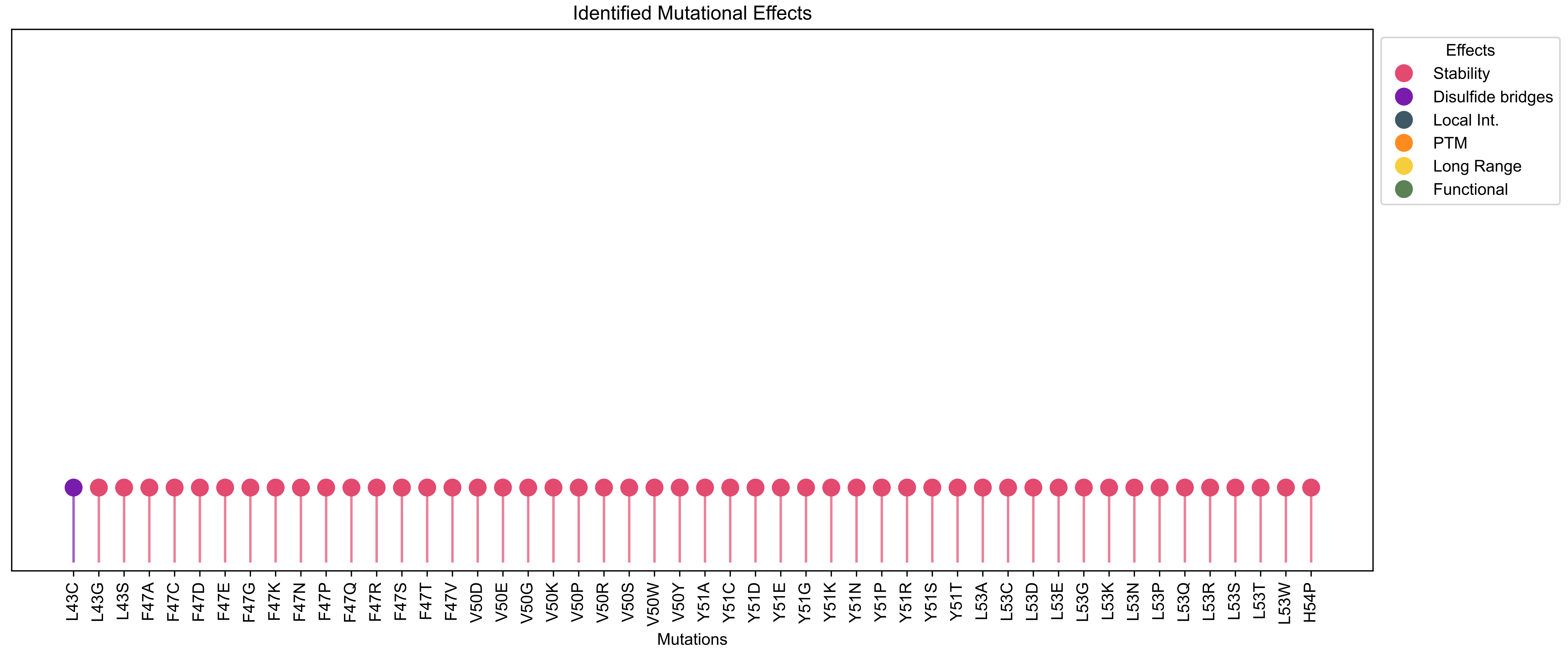

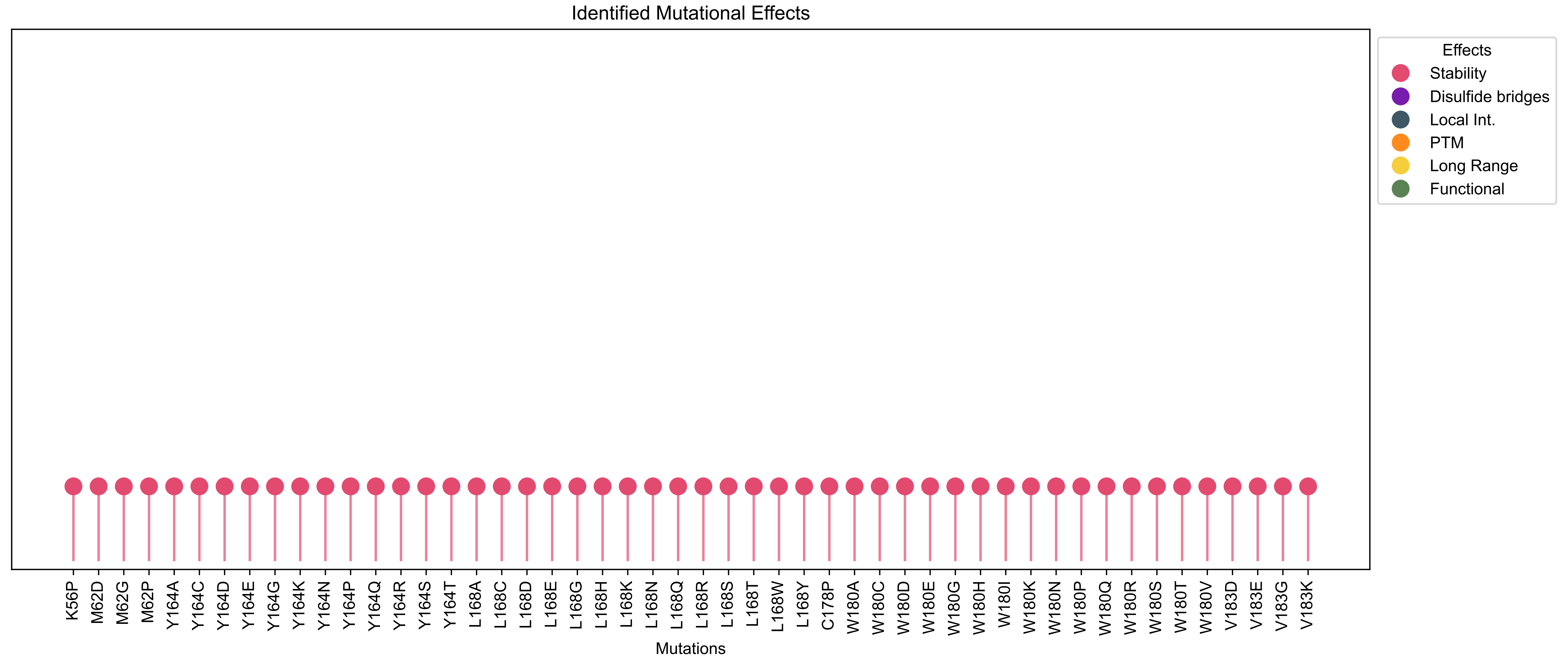

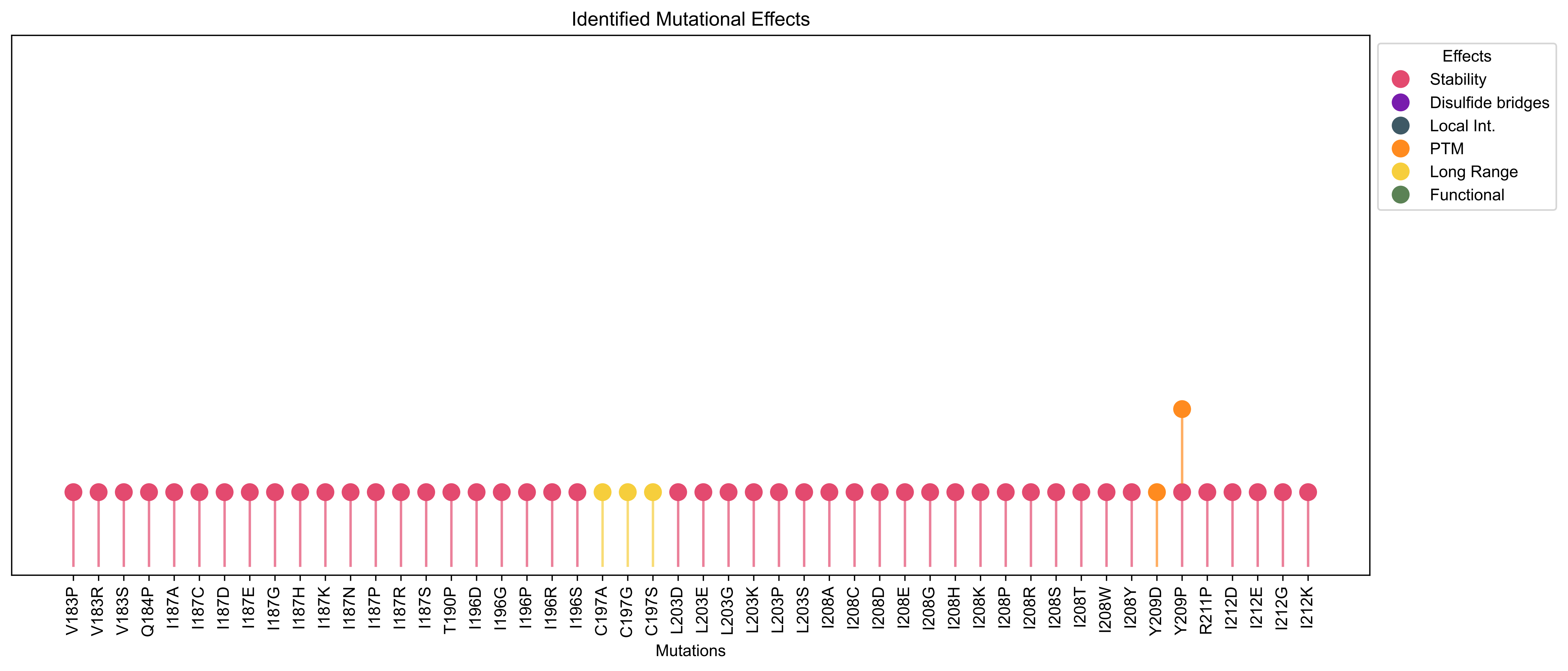

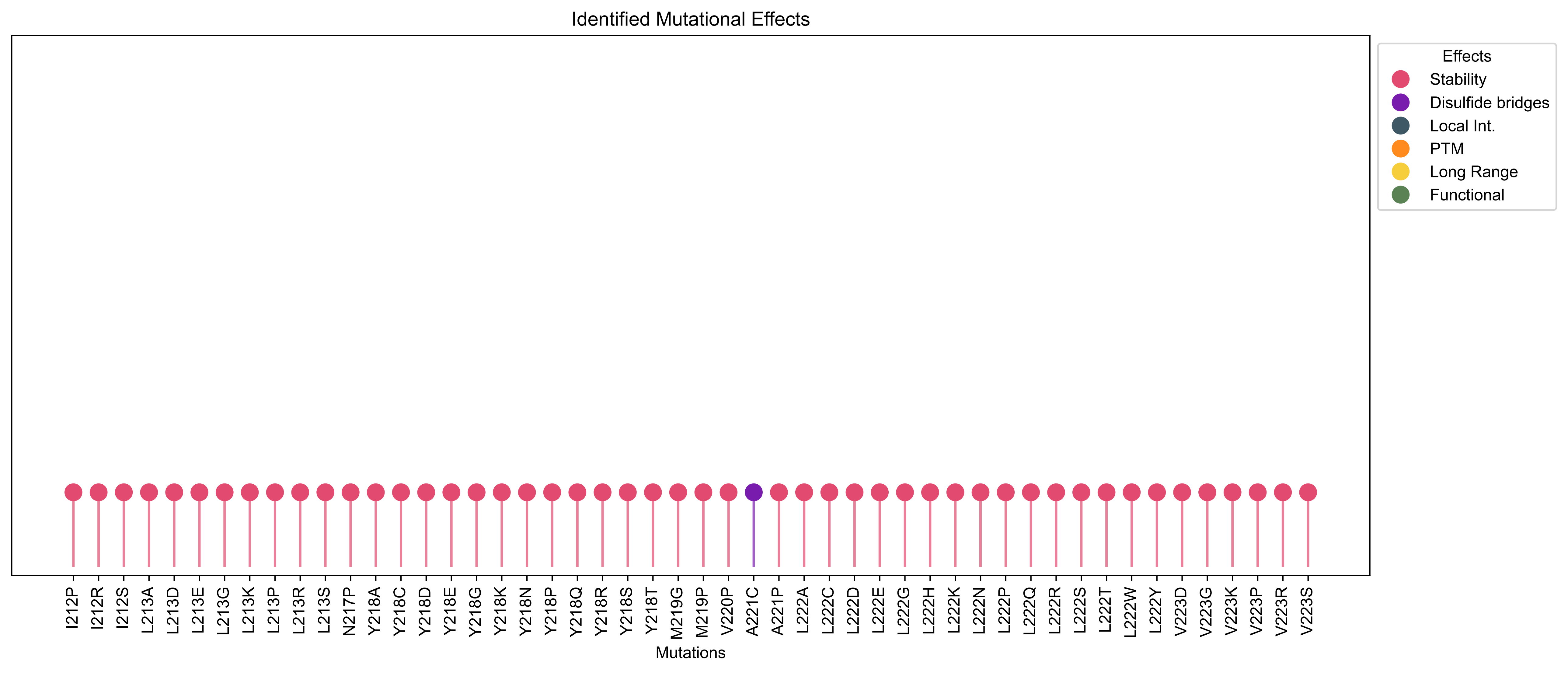

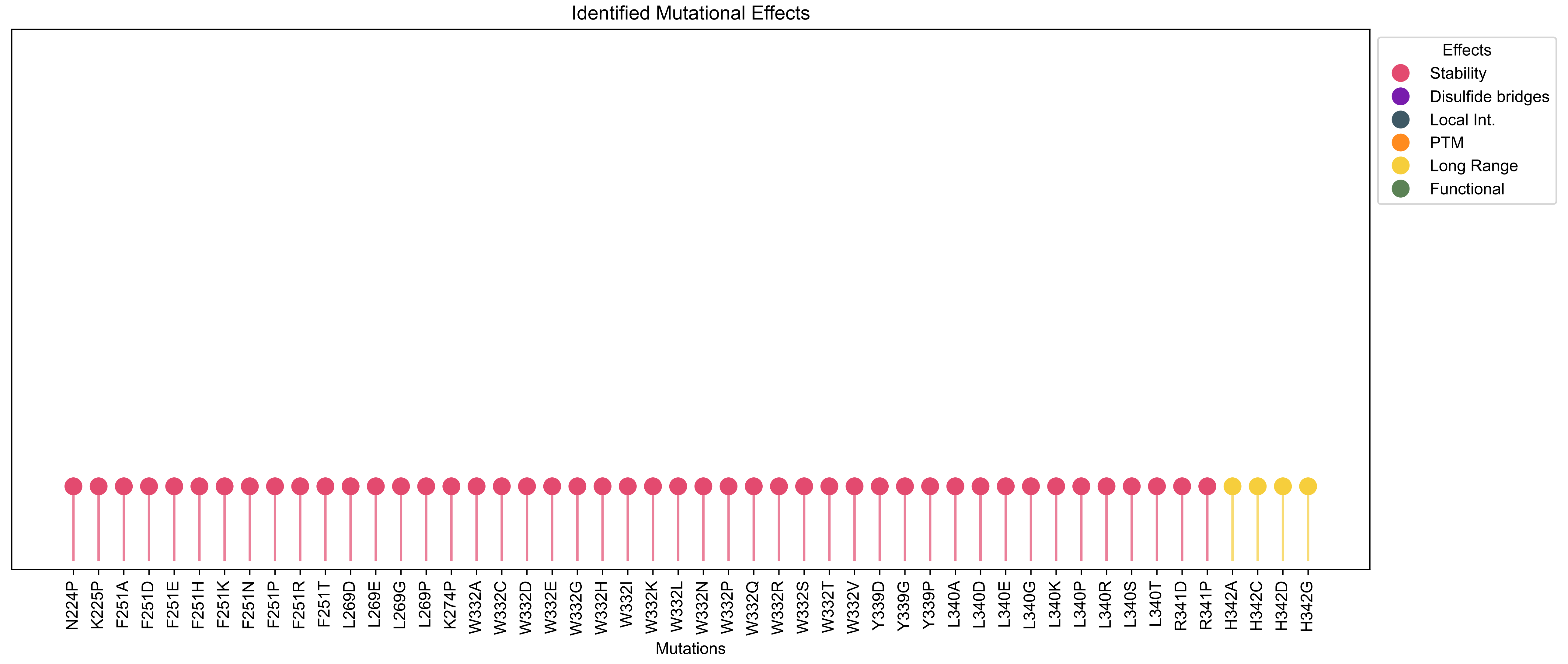

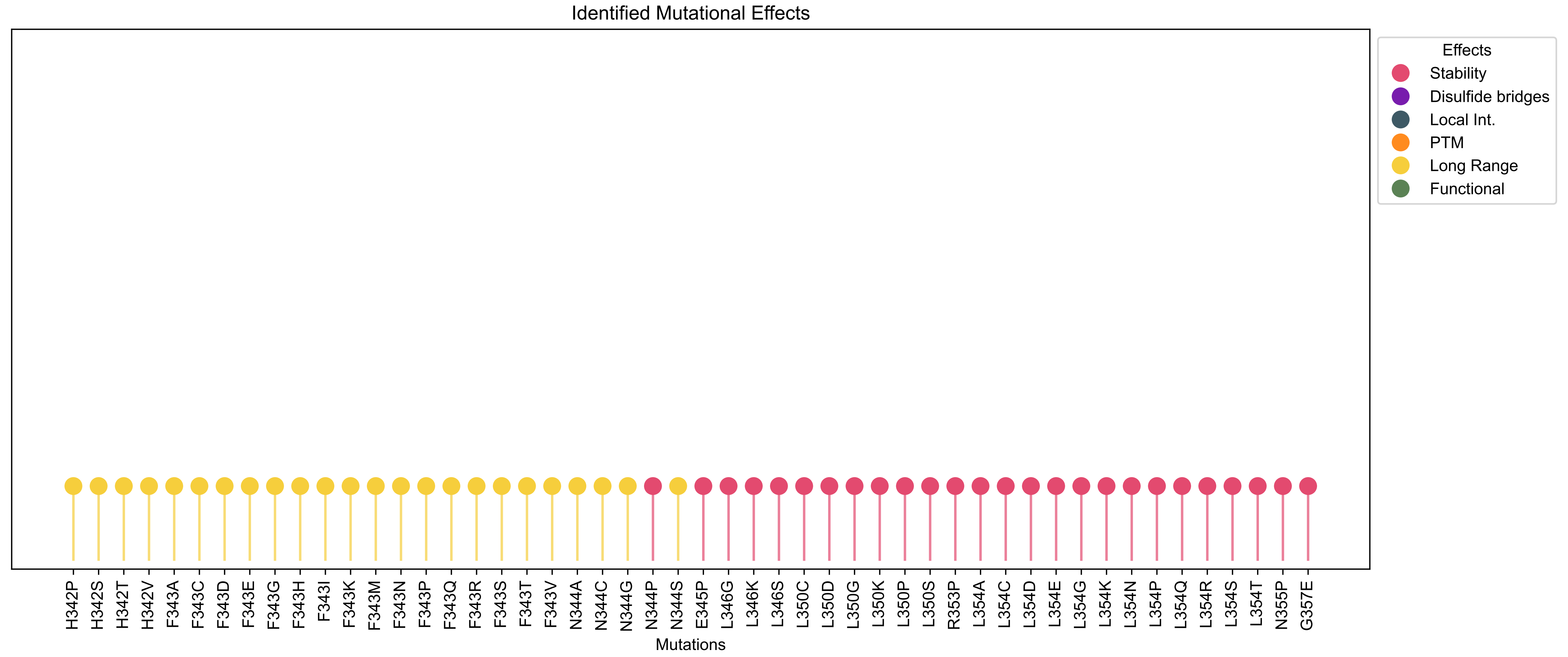

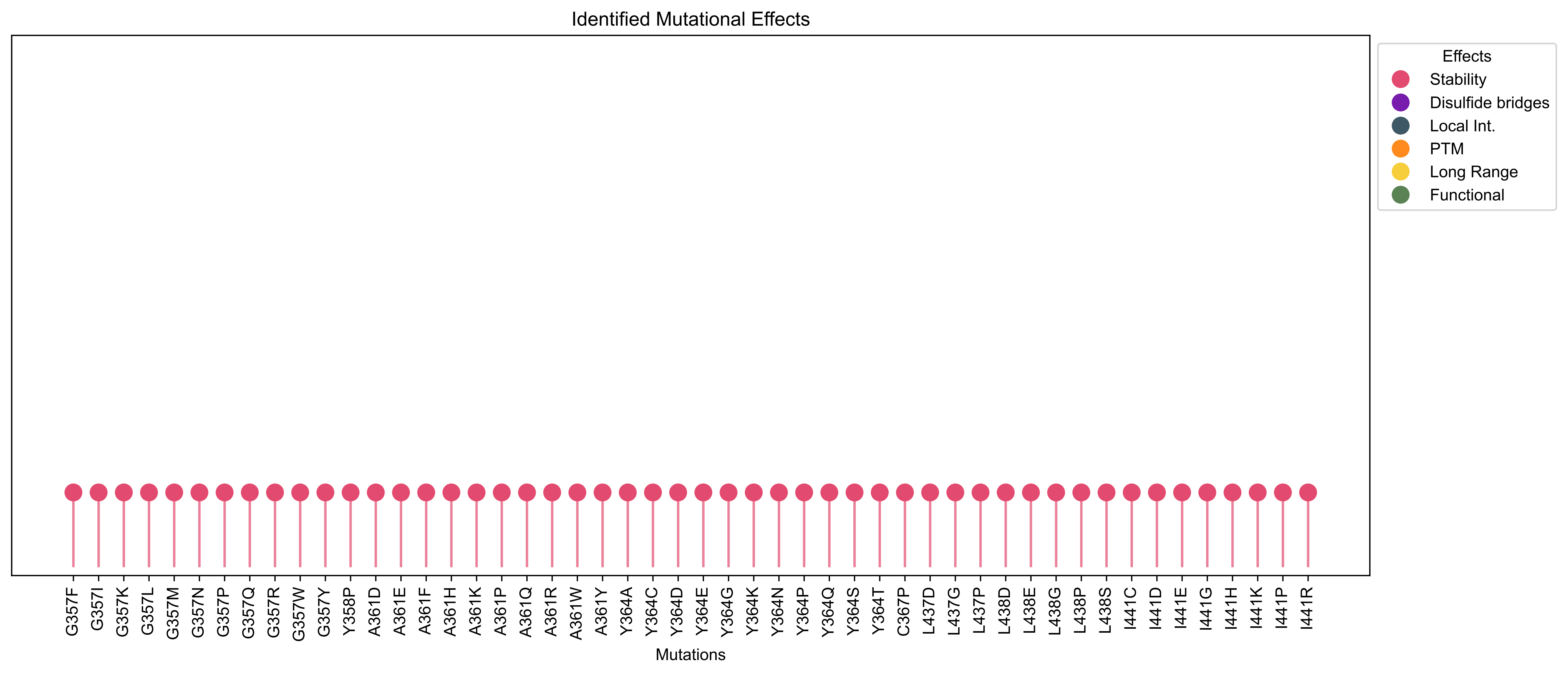

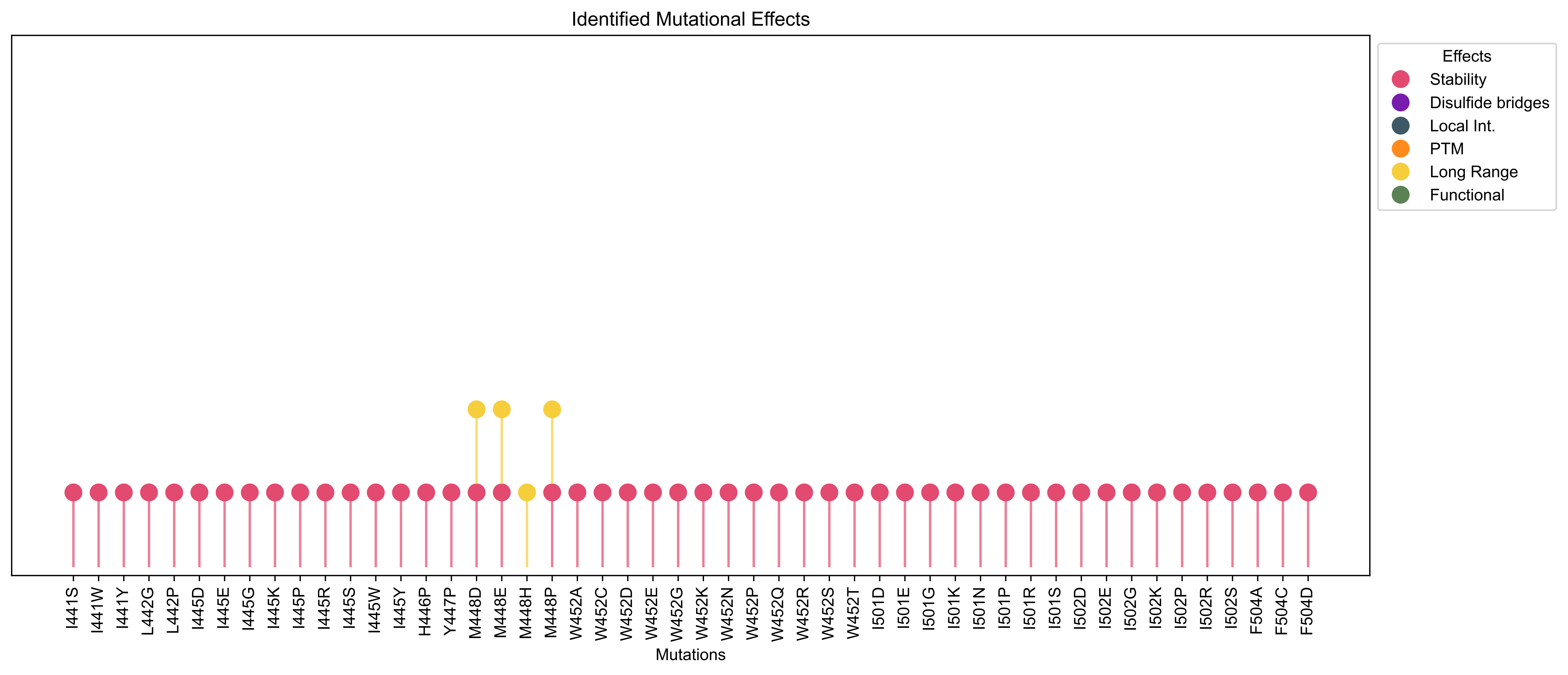

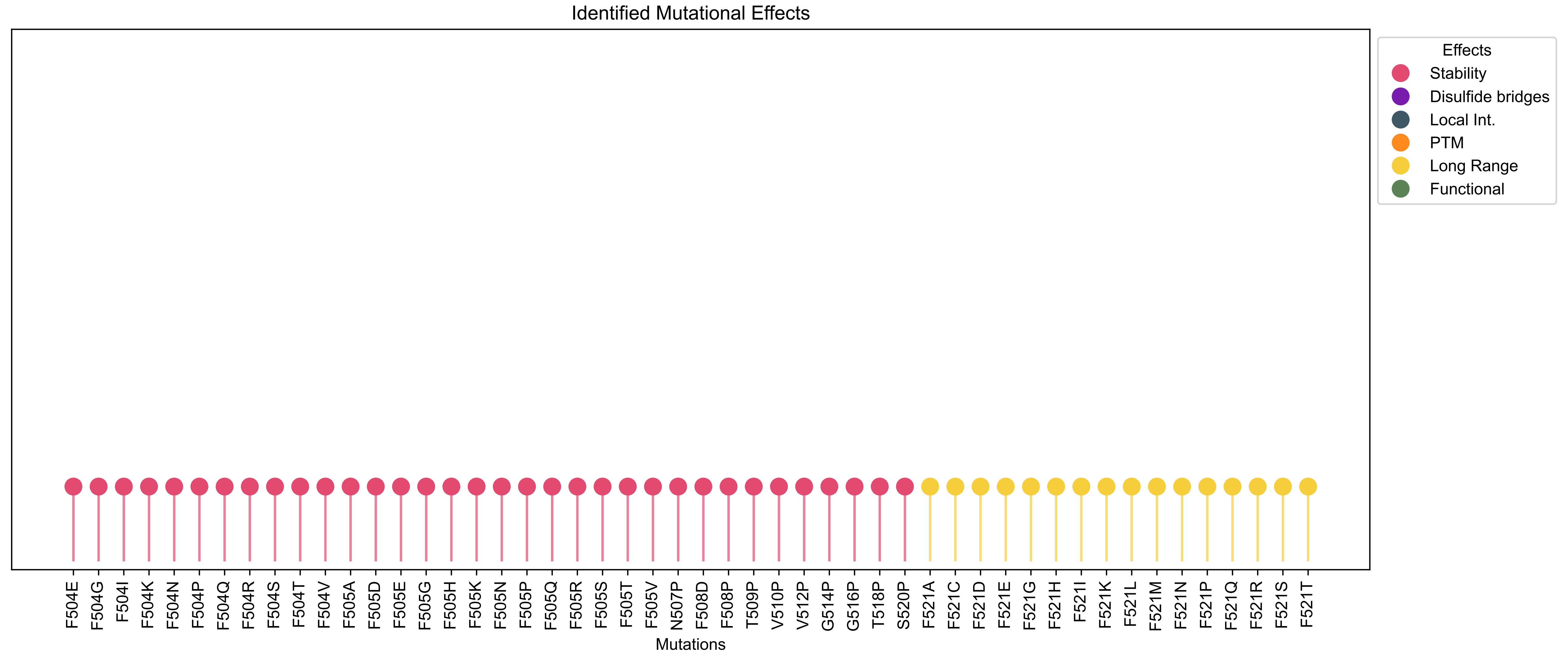

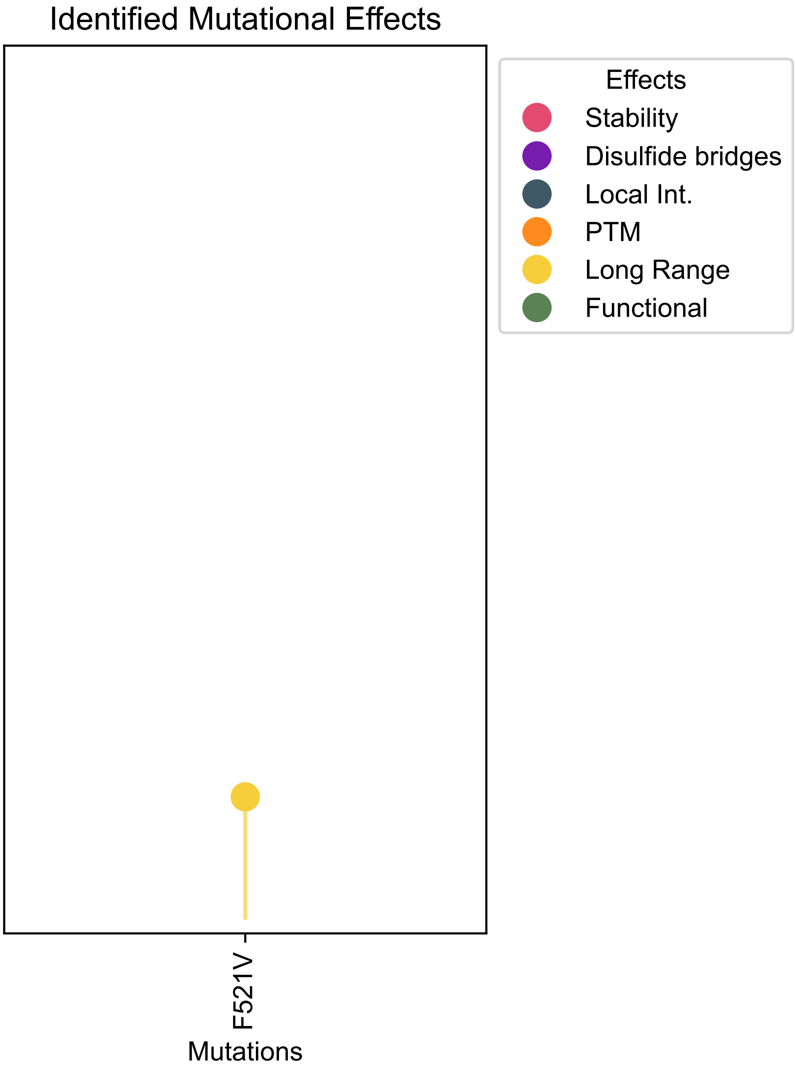

### Figure S3, Lolliplot for the analysis without the DeMaSk filtering

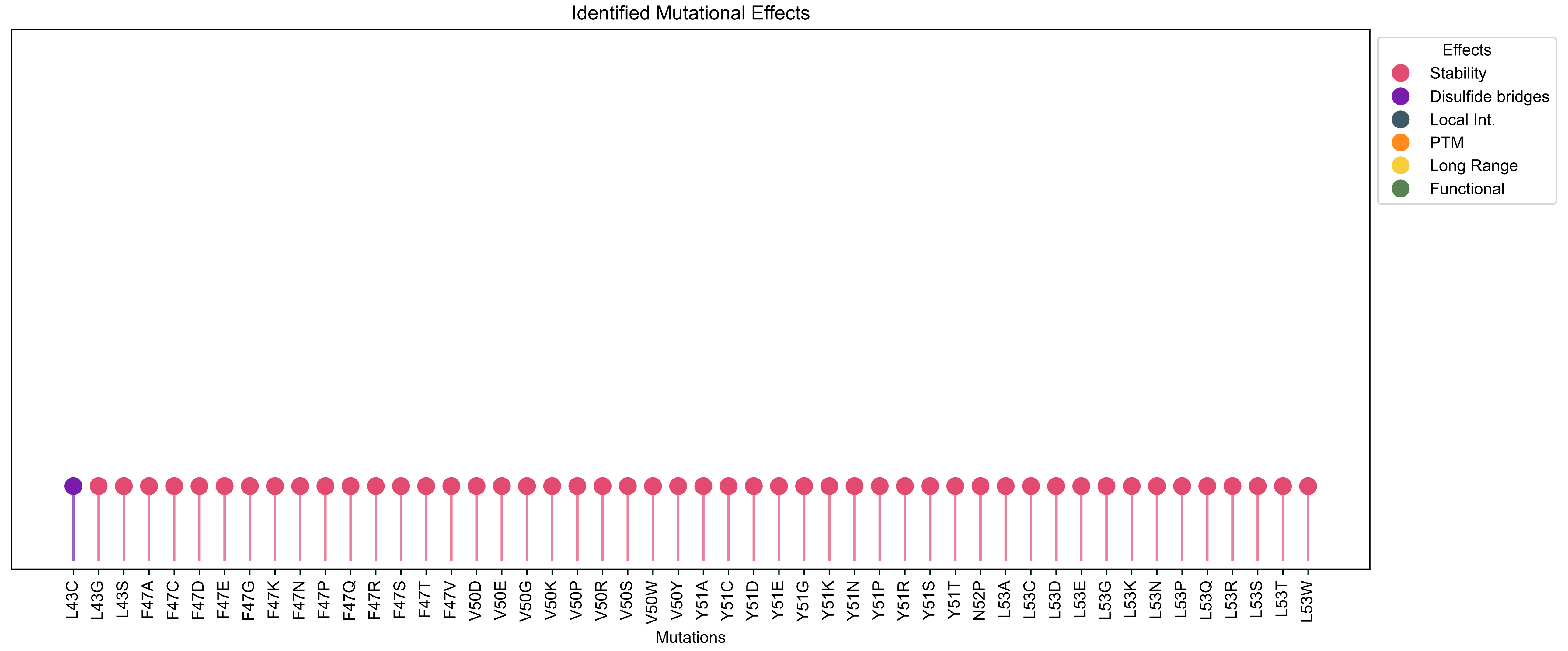

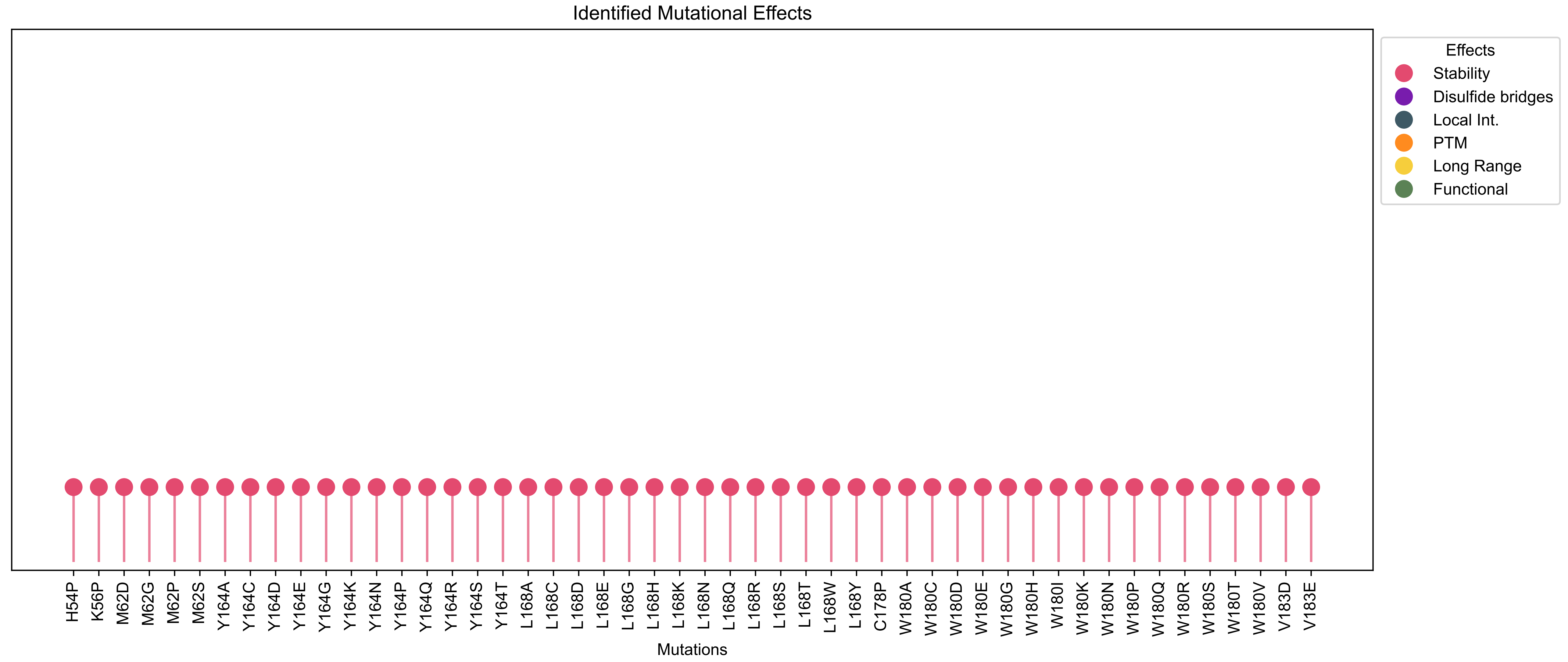

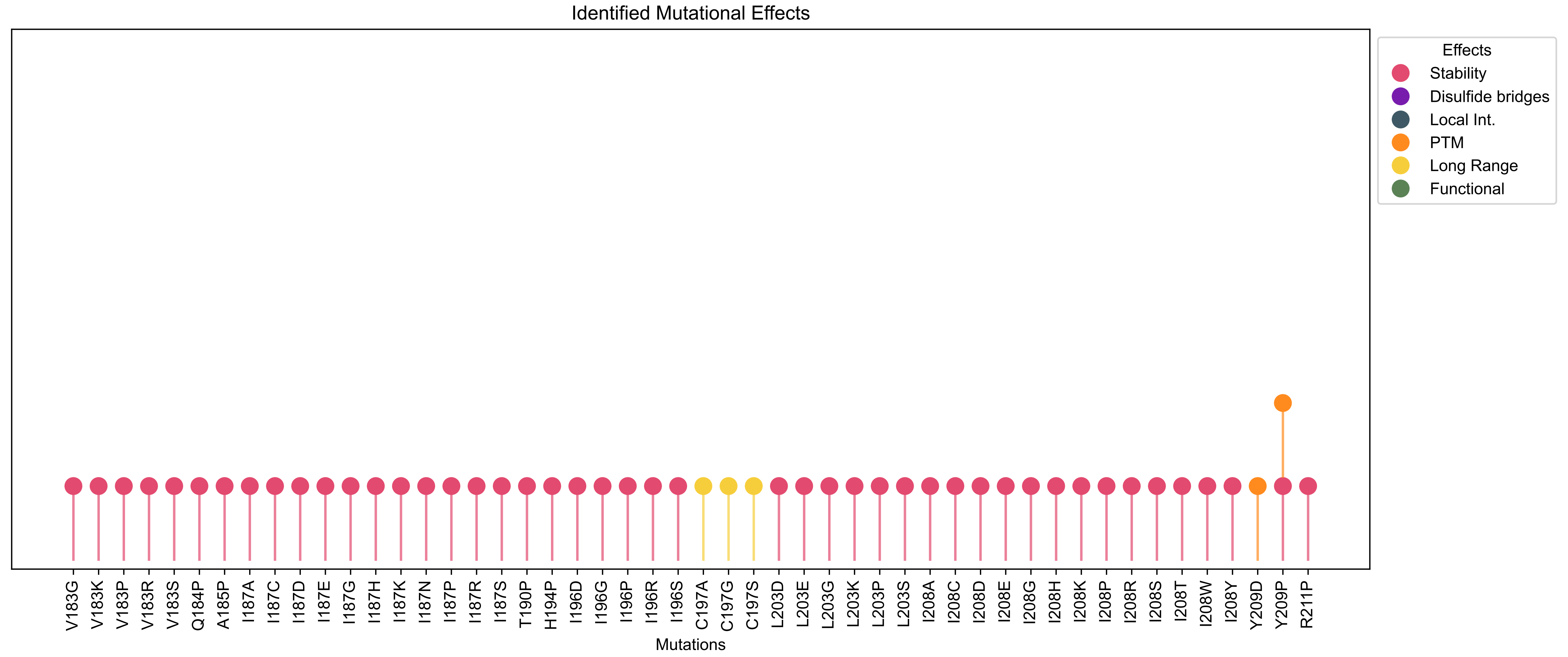

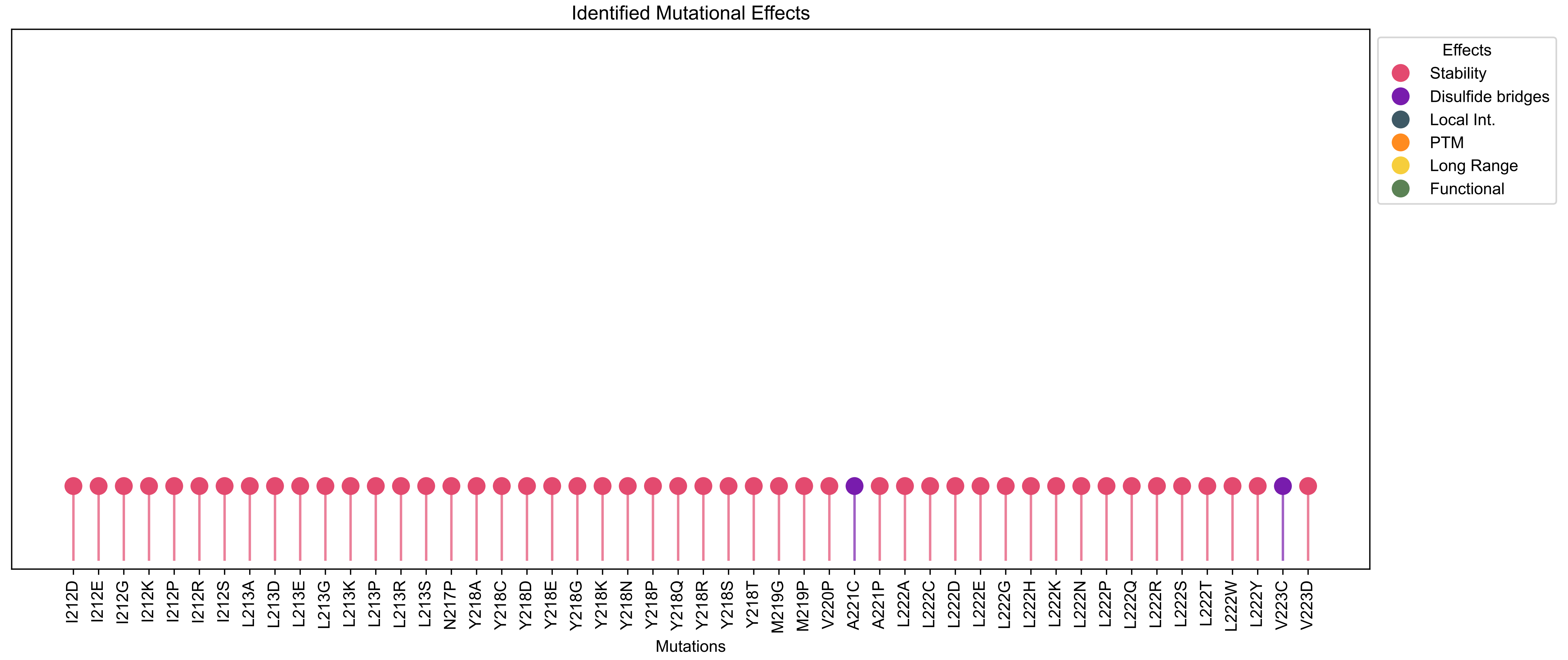

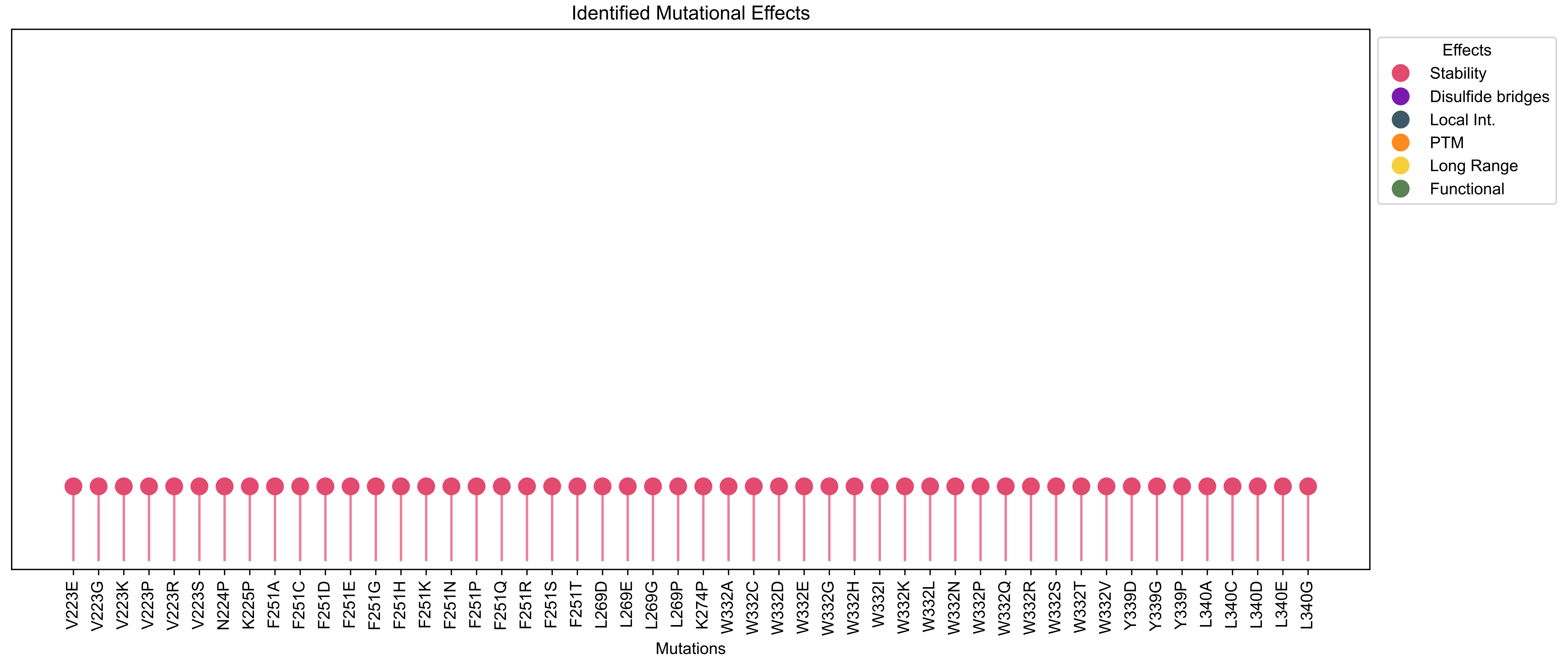

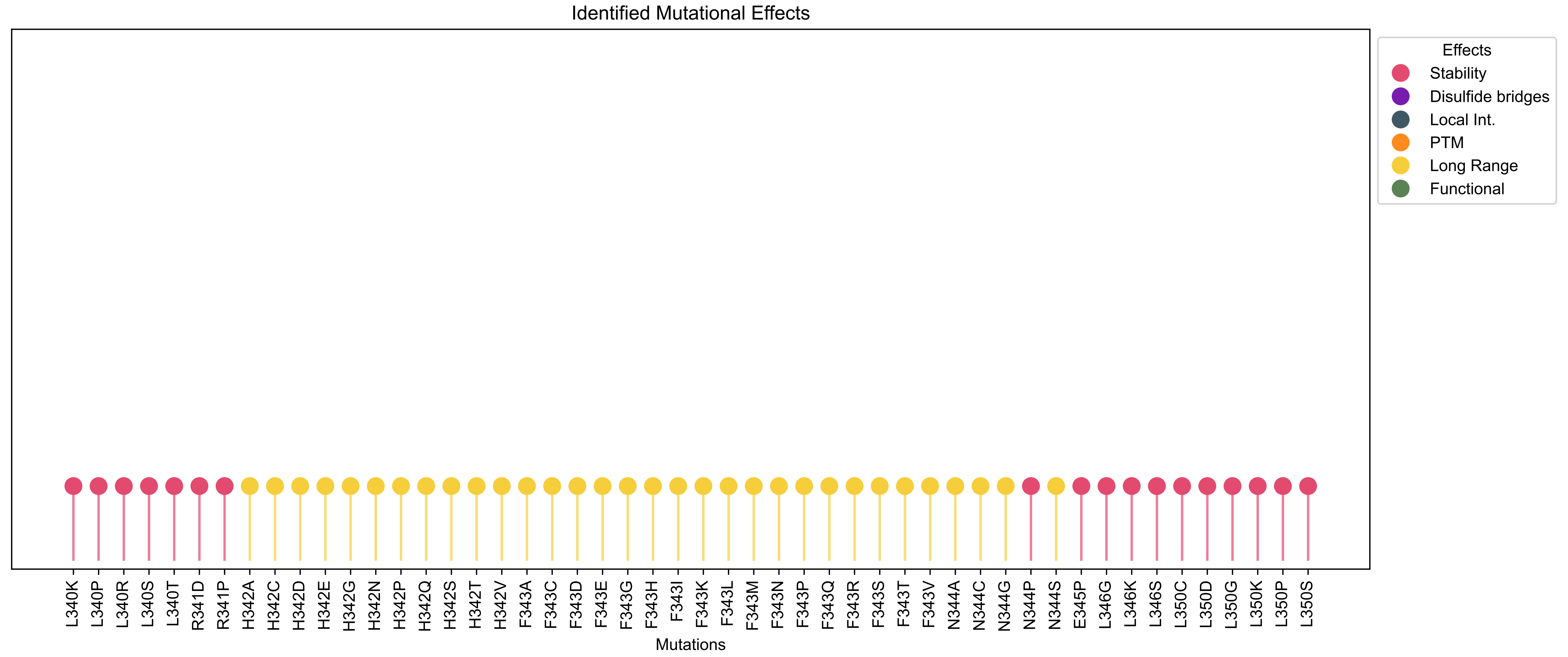

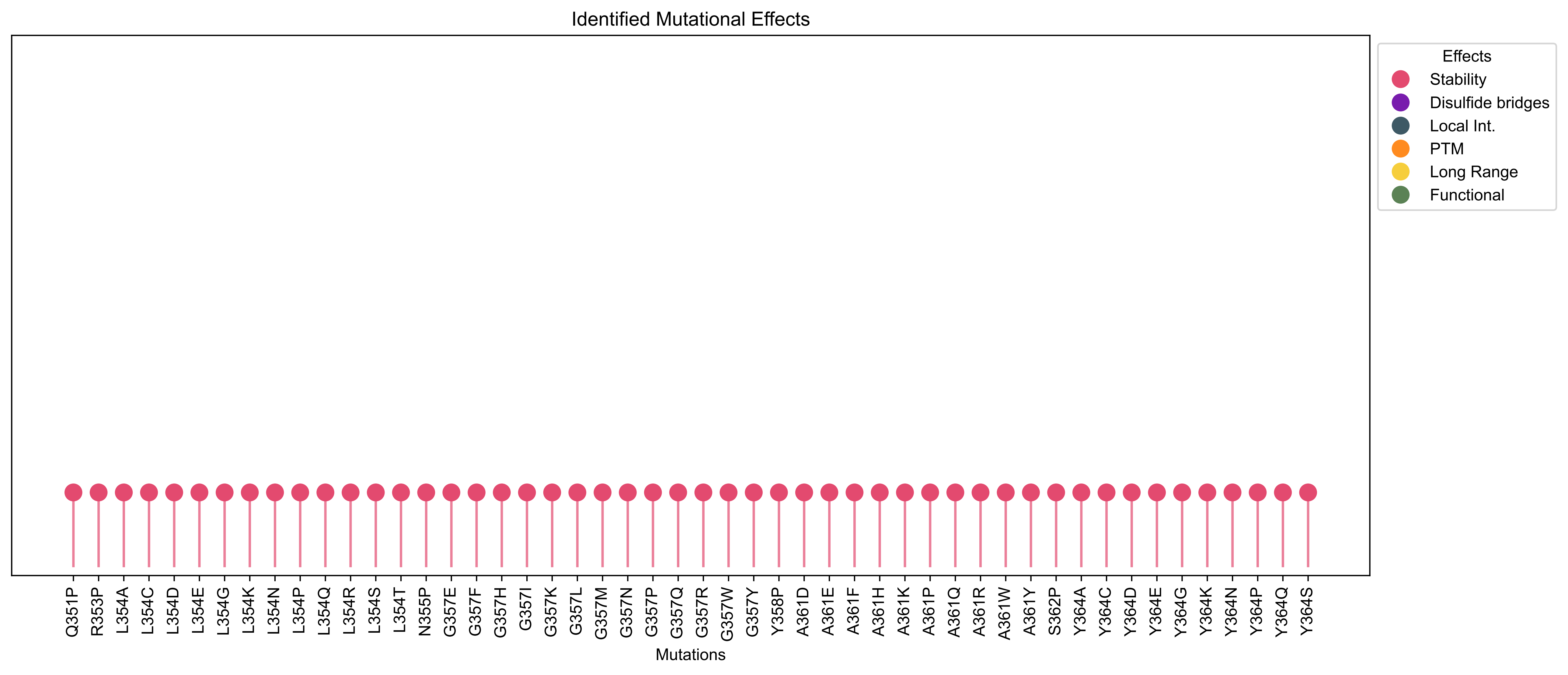

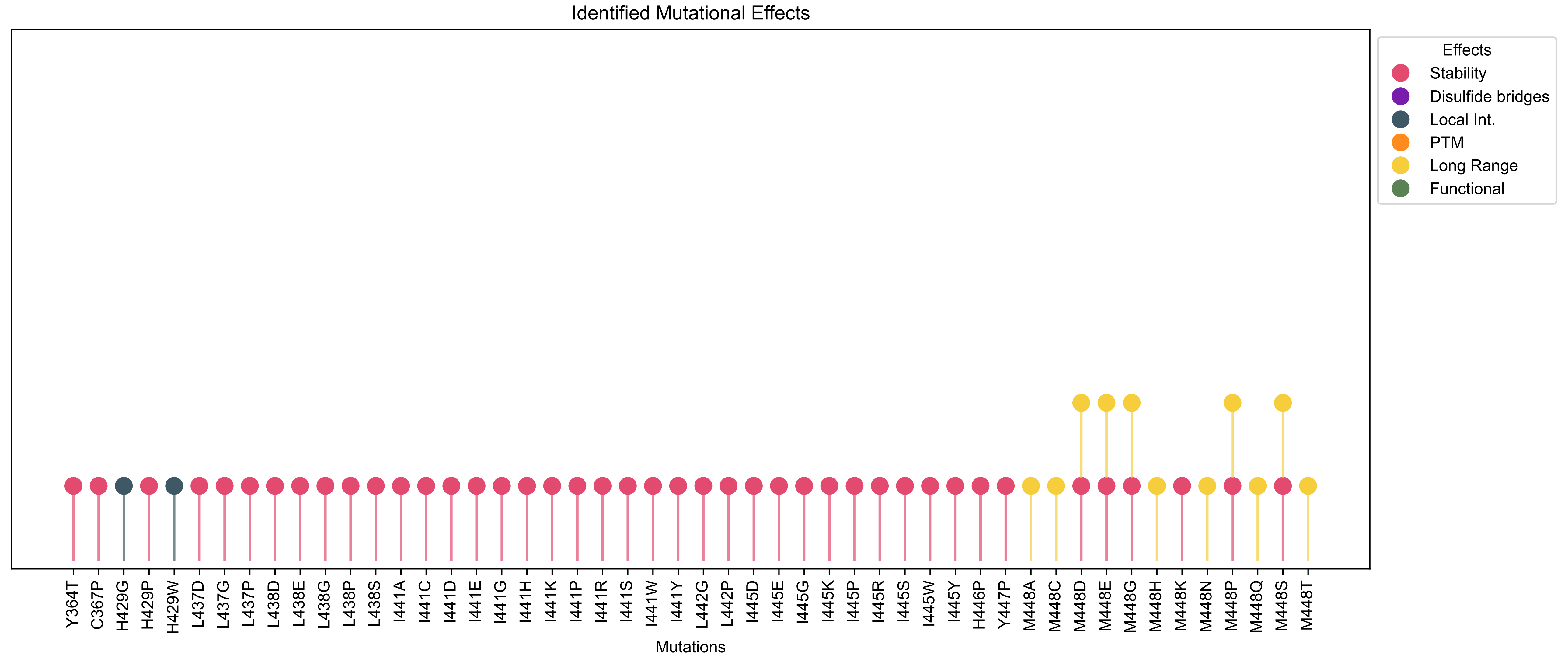

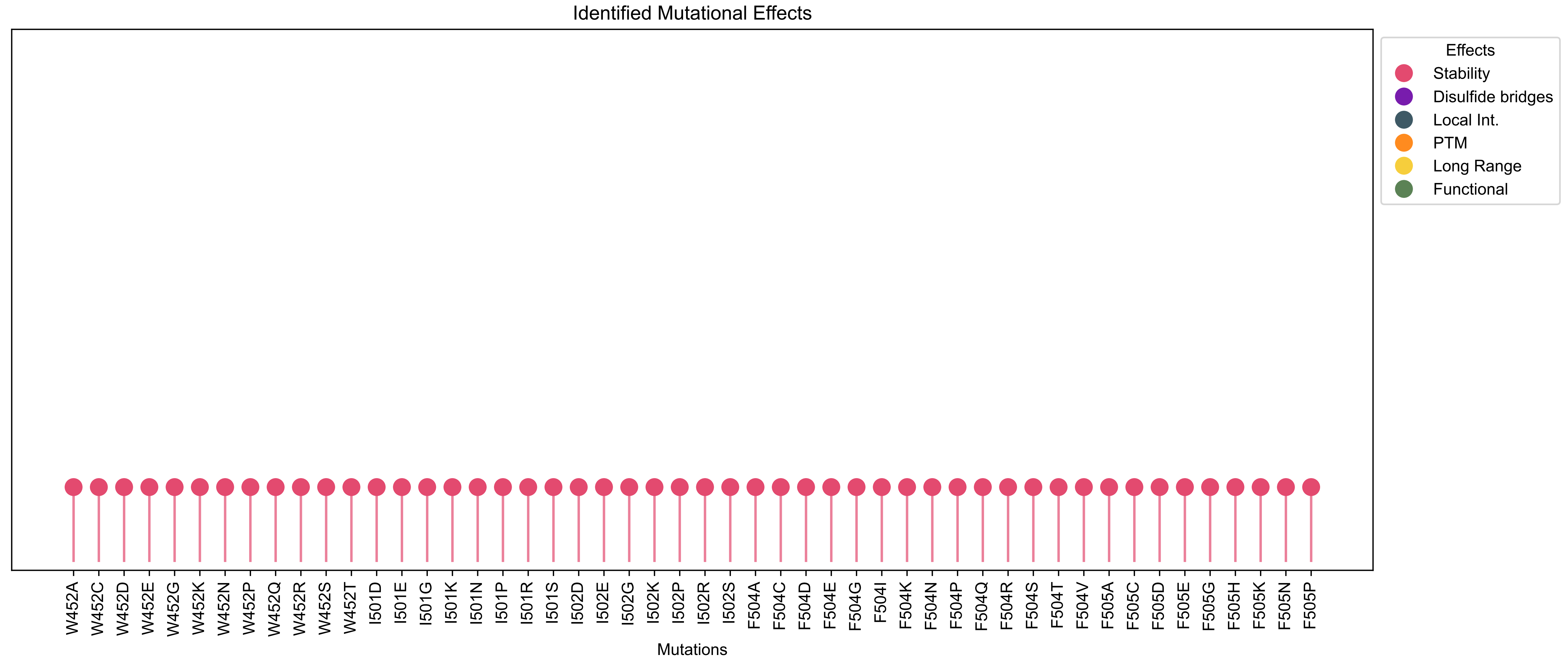

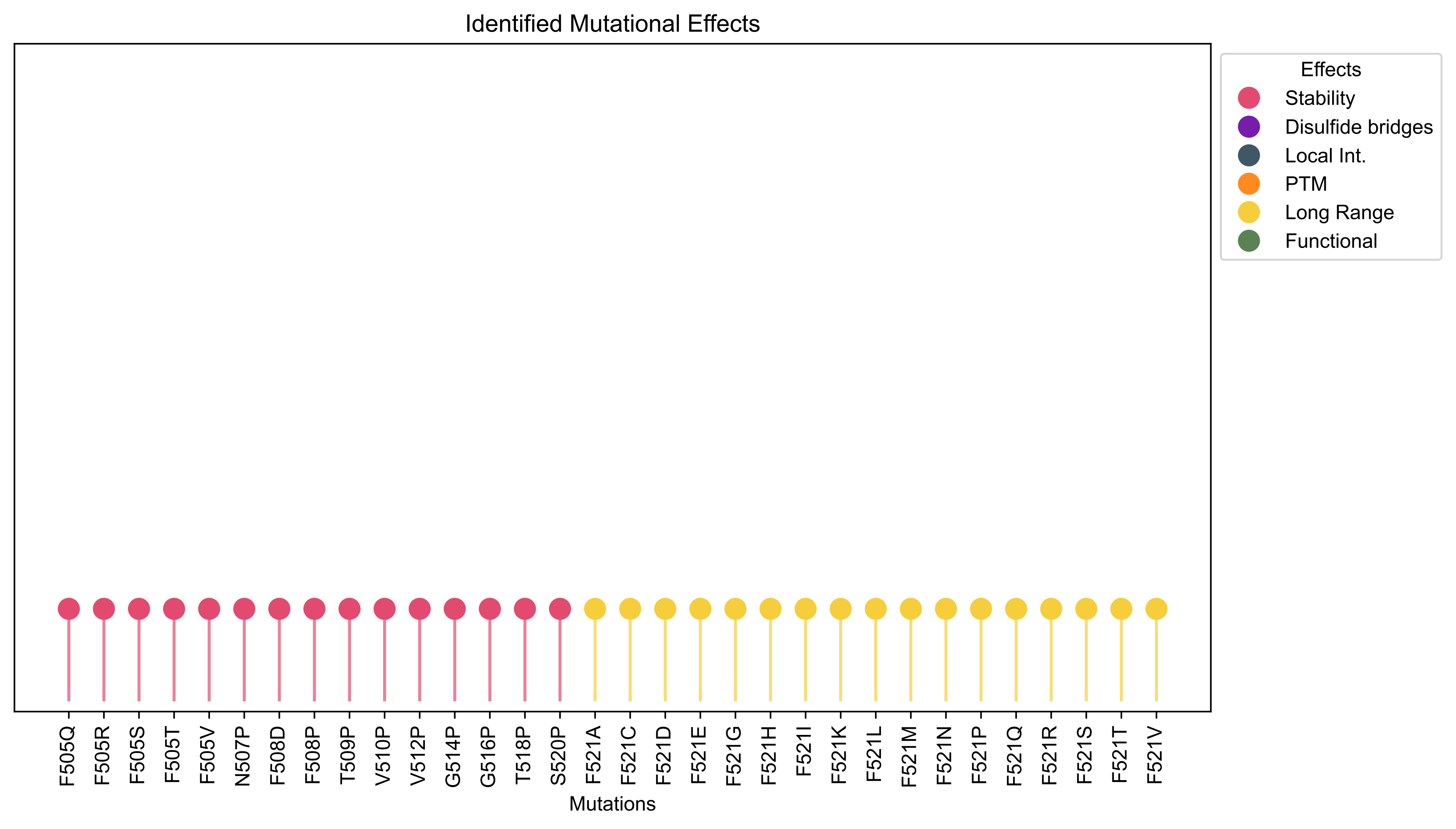
